# New C3H *Kit*^N824K/WT^ cancer mouse model develops late-onset malignant mammary tumors with high penetrance

**DOI:** 10.1101/2022.08.23.504938

**Authors:** Tanja Klein-Rodewald, Kateryna Micklich, Adrián Sanz-Moreno, Julia Calzada-Wack, Monica Tost, Thure Adler, Matthias Klaften, Sibylle Sabrautzki, Bernhard Aigner, Markus Kraiger, Valerie Gailus-Durner, Helmut Fuchs, German Mouse Clinic Consortium, Albert Gründer, Heike Pahl, Eckhard Wolf, Martin Hrabe de Angelis, Birgit Rathkolb

**Affiliations:** Institute of Experimental Genetics, German Mouse Clinic, Helmholtz Zentrum München, German Research Center for Environmental Health, Neuherberg, Germany; Research Unit Comparative Medicine, Helmholtz Zentrum München, German Research Center for Environmental Health, Neuherberg, Germany; Institute of Molecular Animal Breeding and Biotechnology, Gene Center, Ludwig-Maximilians-University München, Munich, Germany; Section of Molecular Hematology, Department of Hematology/Oncology, Universitäts Klinikum Freiburg, Freiburg, Germany; German Center for Diabetes Research (DZD), Neuherberg, Germany; Chair of Experimental Genetics, TUM School of Life Sciences, Technische Universität München, Freising, Germany

**Keywords:** KIT oncogene, mouse model, N824K mutation, N822K mutation, breast cancer, GIST, polycytemia

## Abstract

Gastro-intestinal stromal tumors and acute myeloid leukemia induced by activating stem cell factor receptor tyrosine kinase (KIT) mutations are highly malignant. Less clear is the role of KIT-mutations in the context of breast cancer. Treatment success of KIT-induced cancers is still unsatisfactory because of primary or secondary resistance to therapy. Mouse models offer essential platforms for studies on molecular disease mechanisms in basic cancer research. In the course of the Munich N-ethyl-N-nitrosourea (ENU) mutagenesis program a mouse line with inherited polycythemia was established. It carries a base-pair exchange in the Kit gene leading to an amino acid exchange at position 824 in the activation loop of KIT. This KIT-variant corresponds to the N822K mutation found in human cancers, which is associated with imatinib-resistance. C3H KitN824K/WT mice develop hyperplasia of interstitial cells of Cajal and retention of ingesta in the cecum. In contrast to previous KIT-mutant models, we observe a benign course of gastrointestinal pathology associated with prolonged survival. Female mutants develop mammary carcinomas at late onset and subsequent lung metastasis. The disease model complements existing oncology research platforms. It allows for addressing the role of KIT mutations in breast cancer and identifying genetic and environmental modifiers of disease progression.

## Introduction

The proto-oncogene KIT, a cell surface receptor tyrosine kinase, plays an important role in the development of many cell types including erythrocytes, mast cells and interstitial cells of Cajal (ICCs) ^1–3^. Gain-of-function mutations are responsible for neoplasms originating from these cell types. These include gastrointestinal stromal tumors (GIST) originating from ICCs ^4,5^, mastocytosis, acute myeloid leukemia (AML), malignant melanoma and germ cell neoplasms ^6–8^. Further, certain types of malignant breast cancer are associated with KIT overexpression, but the relevance of KIT-mutations for mamma carcinoma development is unclear, since activating KIT-mutations are rarely found in this context ^9,10^. In humans the KIT N822K mutation potentially induces several cancer types, and is associated with resistance to tyrosin kinase inhibitor (TKI) treatment using Imatinib ^11^.

During the past ten years several mouse models carrying activating Kit mutations were established and significantly contributed to the current knowledge about the role of KIT in cancer development ^12–15^. One of the achievements within the Munich ENU mouse mutagenesis program was the establishment of a mouse line (preliminary laboratory name MVD013) representing microcytic polycythemia ^16^. Here we report on its comprehensive characterization concerning hematopoietic abnormalities, development of neoplasms and unexpected side effects. Details on the identification of the causative genetic alteration, a *Kit^N824K^* mutation, which is homologous to KIT N822K mutation of human patients, and a comparison to previously established models are presented. As a consequence of the high conservation of the KIT protein between human and mouse ^17^ the obtained phenotypic data is expected to add valuable information about the role of *KIT* mutations as a primary cause of cancer and other disorders. A potential application is given in the accompanying paper by Kraiger et al. characterizing the disease progression in C3H *Kit*^N824K/WT^ mice across a life span of twelve month by in-vivo magnetic resonance imaging (Kraiger et al., accompanying paper). This novel disease model is expected to facilitate future studies addressing the identification of genetic and environmental factors affecting disease progression in KIT-induced cancer.

## Results

### A heterozygous Kit^N824K^ mutation is associated with polycythemia in MVD013 mice

Microcytic polycythemia in ENU-induced mouse line MVD013 is inherited as a monogenic, dominant trait ^16^. Heterozygous mutant animals show increased red blood cell counts with marked microcytosis (Fig. 1 a-c) resulting in significantly increased hemoglobin (16.8±0.26 g/dl vs. 14.2±0.48 g/dl and 16.7±0.12 g/dl vs. 15.0±0.13 g/dl in males and females respectively, p<0.001) and hematocrit values (60.8±0.82 % vs. 53.2±0.46 % and 59.2±0.35 % vs. 52.1±0.4 % in males and females respectively, p<0.001). Additionally, platelet counts were mildly decreased in mutants compared to controls (Fig. 1d). Although total white blood cell counts were similar in samples collected from mutant and control mice (Fig. 1e), automated leukocyte pre-differentiation and FACS analysis of peripheral blood leukocytes indicated an increased proportion of granulocytic cells (Fig. 1f, Table 1, Supplement 1). Also, MPO-immunohistochemistry revealed an increased proportion of neutrophil granulocytes in spleens of mutant animals (Fig. 1g).

**Figure 1.**
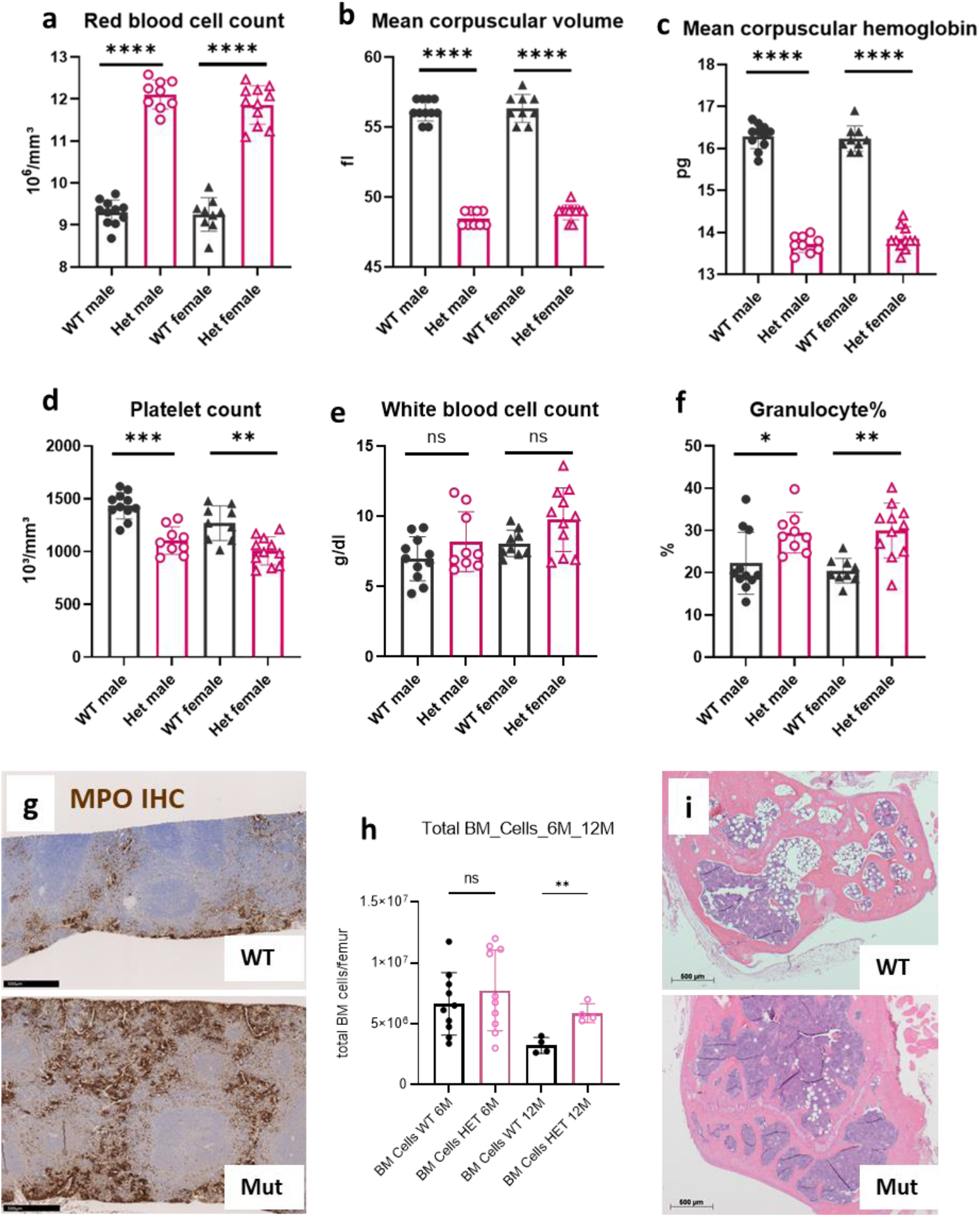
Hematological characterization of C3H *Kit*^N824K/WT^ mutant mice. Red blood cell counts (1a), mean corpuscular volume (1b) and mean corpuscular hemoglobin (1c), platelet counts (1d), total white blood cell counts (1e) and percentage of granulocytic cells (1f) of controls (WT) and heterozygous (Het) mutant animals (n= 9-11 per group, age 20-21weeks), representative pictures of IHC for MPO of the spleen of a control (upper panel) and a mutant animal (lower panel) at age 6 months (1g), Cell counts of femoral bone marrow derived from female control and heterozygous mutant mice collected at age 6 (n=10 per group) or 12 months (n=4 per group) (1h), representative picture of bone marrow histology in the femural epiphysis of 12 months old control (upper panel) and mutant (lower panel) animals (1i). **** p<0.0001; *** p>0.001, ** p<0.01, * p<0.05 (Wilcoxon Rank Sum Test)

**Table 1.** Additional findings in mutants compared to controls from the GMC Primary Phenotyping Screen (age 14 – 22 weeks).

| Screen | Test | Summary of results |
| --- | --- | --- |
| Dysmorphology | Anatomical observation, DEXA, X-ray | No clear genotype-related effects on outer appearance, decreased BMC, BMD and body fat proportion in female mutants |
| Behaviour | Open Field, PPI | No effects on Open Field behavior, Startle Response or Prepulse Inhibition observed |
| Neurology | SHIRPA, Grip Strength, Rotarod | No differences between groups observed during SHIRPA protocol, no differences in grip strength, slightly improved performance on the rotarod |
| Eye | Slit lamp, Funduscopy | No differences, between groups but typical findings related to genetic background in mutants and controls |
| Nociception | Hot plate test | No effects on hot plate pain reaction |
| Metabolism | Indirect Calorimetry | No effects on energy turnover or rectal body temperature observed |
| Clinical Chemistry | Glucose Tolerance Test (IpGTT) | Decreased basal fasting glucose level in male mutants, improved glucose tolerance in both sexes |
|  | Clinical Chemistry (fasting) | Decreased fasting glucose, cholesterol and triglyceride levels |
|  | Clinical Chemistry (fed) | Decreased plasma transferrin and iron levels as well as plasma total protein and albumin concentrations, elevated potassium concentration and ASAT activity. |
| Cardiovascular Screen | Blood pressure, Echocardiography, Heart weight | Decreased diastolic blood pressure in female mutants, slightly decreased heart rate in male mutants, slightly decreased heart weight corresponding to decreased body weight in mutants |
| Lung function | Whole body plethysmography | Decreased breathing frequency of female mutants during sleep. |
| Immunology | FACS Analysis of blood leukocytes, Immunoglobulins | Mainly in females elevated proportion of granulocytes and decreased proportion of lymphocytes in peripheral blood; Increased plasma concentrations of IgM, IgA, IgG3, IgG2a and IgG2b |
| Steroid Screen | DHEA, testosterone | Decreased DHEA in female mutants |

Abbreviations: DEXA – Dual-energy X-ray absorptiometry, BMC – bone mineral content, BMD – bone mineral density, PPI – Pre pulse inhibition, SHIRPA – Smithkline Beecham Harwell Imperial College phenotype assessment, ip – intra peritoneal, FACS – fluorescence associated cell sorting, ASAT – aspartate aminotransferase, Ig – Immunoglobuline, DHEA – dehydroepiandrosterone.

Bone marrow total cell counts were similar in six months old mutant and control females, but significantly elevated compared to controls at 12 months in mutants (Fig. 1h). In line with this more hematopoietic and fewer fat cells were observed histologically in femoral bone marrow of one year old mutant animals (Fig. 1i). Colony assays revealed increased proliferative activity of stem cells from younger heterozygous mice, but decreased proliferation and elevated apoptosis rates of stem cells from older mutants (Supplement 2, Fig. S1).

A classical linkage analysis based on genotyping of 161 single nucleotide polymorphisms (SNPs) evenly distributed on chromosomes 1-19 was carried out in 61 N2-hybrid mice including 23 animals showing polycythemia and 38 littermates showing a wild-type phenotype. The strongest linkage of polycythemia with C3B6 heterozygosity was observed for two SNPs on chromosome 5: rs29635956 at about 67.73 mbp and rs31585424 at about 79.10 mbp. No other chromosomal region showed remarkable linkage of the phenotype to C3B6 heterozygosity.

Exome sequencing identified three heterozygous mutations affecting amino-acid sequence in the respective gene products, which were present in the exomes of the two mutant MVD013 mice but not in any wild-type or mutant mouse from other lines on C3H background: A missense mutation in *Ifi203* on chromosome 1, a missense mutation in *Kit* on chromosome 5 and a missense mutation in *Myh13* on chromosome 11. Only the *Kit* mutation, a T to A substitution in exon 17, was localized within the identified mapping region on chromosome 5. It was confirmed by sequencing of the respective region in two additional animals with mutant and control phenotype (Fig. 2a). According to international rules of nomenclature the mutant line was renamed C3H *Kit*^N824K/WT^, since the mutation results in the replacement of an Asparagine residue by Lysine in amino acid position 824. Alignment of the mouse and human Kit amino acid sequences reveals that the mutation is homologous to the N822K mutation found in human patients.

**Figure 2.**
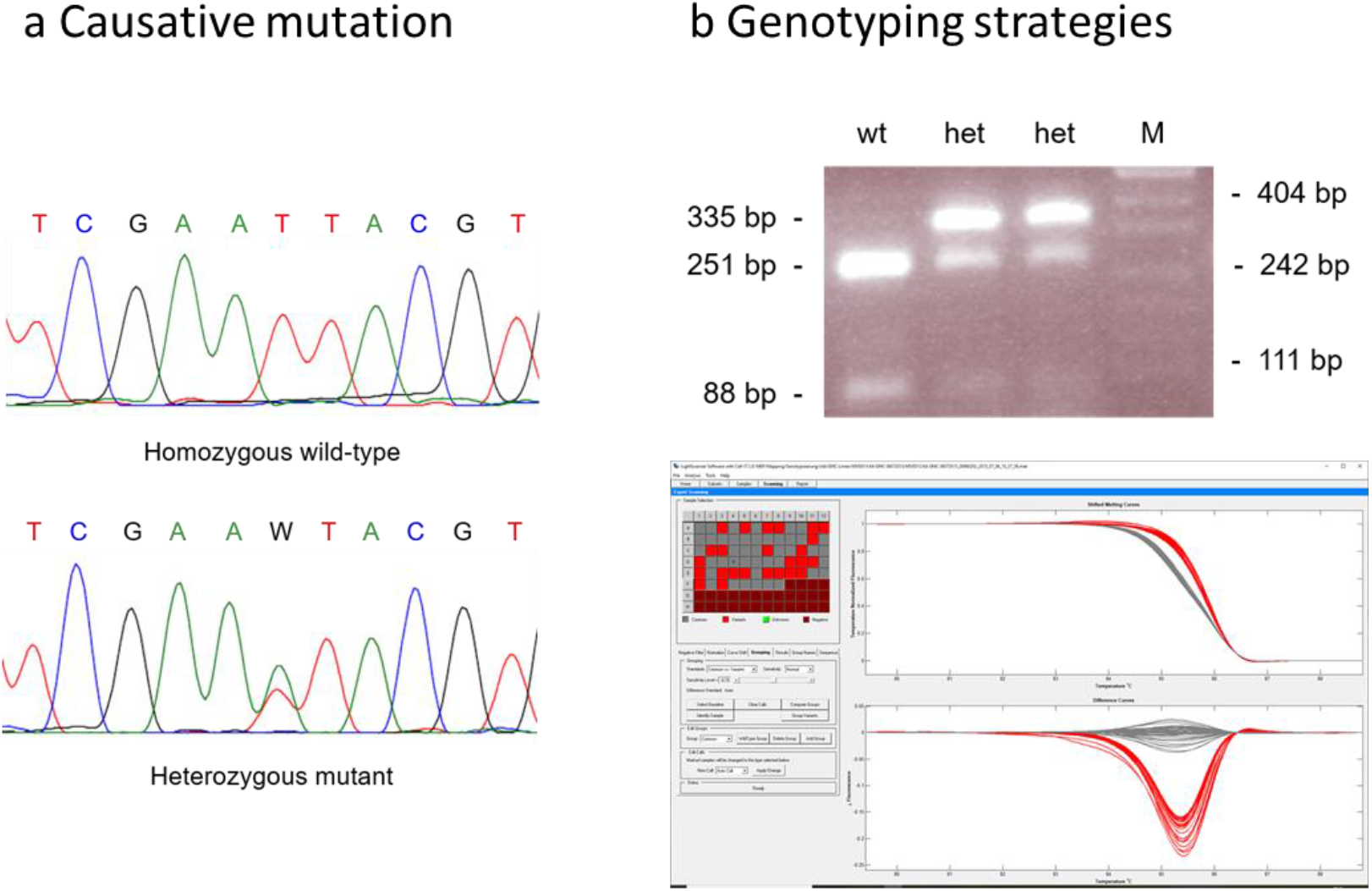
Genetic characterization of C3H *Kit*^N824K/WT^ mutant mice: Sequence tracing of the mutation in exome 17 of the *Kit* gene (2a): sequence obtained from a control animal (upper panel) and from heterozygous animal showing the mutant phenotype (lower panel), (2b): exemplary depiction of genotyping results using PCR-RFLP method (upper panel) or heteroduplex detection using the lightscanner device (lower panel).

Genotyping procedures based on PCR with restriction enzyme fragment length polymorphism (PCR-RFLP, Fig. 2b upper panel) or PCR followed by heteroduplex detection (Fig. 2b lower panel) were established. In 76 animals classified by PCR-RFLP and CBC genotypic and phenotypic classification as mutant or wild-type matched for all but 2 animals. In 53 animals subjected to blood cell count analysis and genotyping by heteroduplex analysis, the outcomes for one animal did not fit together. These results demonstrate an almost complete match of the mutation with the aberrant phenotype and indicate, that CBC phenotyping and genotyping yield similar accuracy concerning correct classification of mutant animals and wild-type littermates in this line.

### Polycythemia in Kit^N824K/WT^ mutant mice is fully penetrant and homozygosity is most likely lethal

Microcytic polycythemia in mouse line MVD013 is inherited as a monogenic, dominant trait, with the expected proportion (50.1% mutant vs. 49.9% WT) of offspring derived from matings of heterozygous mutant mice to wild-type mates showing the mutant phenotype. Together with the almost complete genotype-phenotype match for animals classified by CBC and genotyping these results indicate, that polycythemia is fully penetrant in heterozygous mutant animals. Polycythemic progeny in litters derived from mating heterozygous male and female animals (57.5% mutant and 42.5% WT), however, did not reach the proportion of around 75% of offspring with mutant phenotype expected in case of full viability of homozygous and heterozygous mutant offspring. The proportion of mutant animals was even lower than 66% that would have been expected according to Mendel in case of full viability of heterozygous offspring and complete pre- or perinatal lethality of homozygous animals. This observation suggests homozygous lethality and reduced viability of heterozygous offspring when derived from heterozygous parents. The assumption of homozygous lethality is also supported by the finding of a decreased mean litter size with in average 4.5 living offspring per litter instead of 5.2 in C3H-WT maintenance breeding. However, since litters derived from heterozygous parents mated to wild-type partners were rather bigger than mean litter size obtained from C3H maintenance breeding (6.1 born, 5.8 weaned per litter vs. 5.4 born, 5.2 weaned in C3H), fertility of heterozygous mutants in general appeared unaffected.

### Mutant animals develop hyperplasia of ICCs and retention of ingesta in the cecum

Macroscopic and histological evaluation of 76 mutant animals ranging from age less than 2 months up to 20 months revealed signs of abnormal cecum morphology and altered intestinal passage.

Hyperplasia of ICCs in the myenteric plexus region was detected in all male and female *Kit* mutants, older than 2 months, which equates to 100 % penetrance of this gastrointestinal pathology. Except two very young female mutant mice (aged 2 months) all mutants displayed diffuse ICC hyperplasia in the stomach involving the complete glandular region and the limiting ridge separating glandular and forestomach (Fig. 3 a-c). In around 50% of the animals, independent of age and sex, also the muscularis of the glandular stomach was affected (Fig.3 d-i). Immunohistochemistry revealed KIT-positivity of the hyperplastic cells (Fig. 3 j-l). Additional hyperplastic regions were found in cecum and colon of all but two mutant mice (Fig. 3, m, n, q), while the small intestine was not affected. The hyperplastic cells displayed a spindle-shaped morphology with very low if any mitotic activity. Thickness of the hyperproliferative layer was only slightly increased in old compared to young mutants.

**Figure 3.**
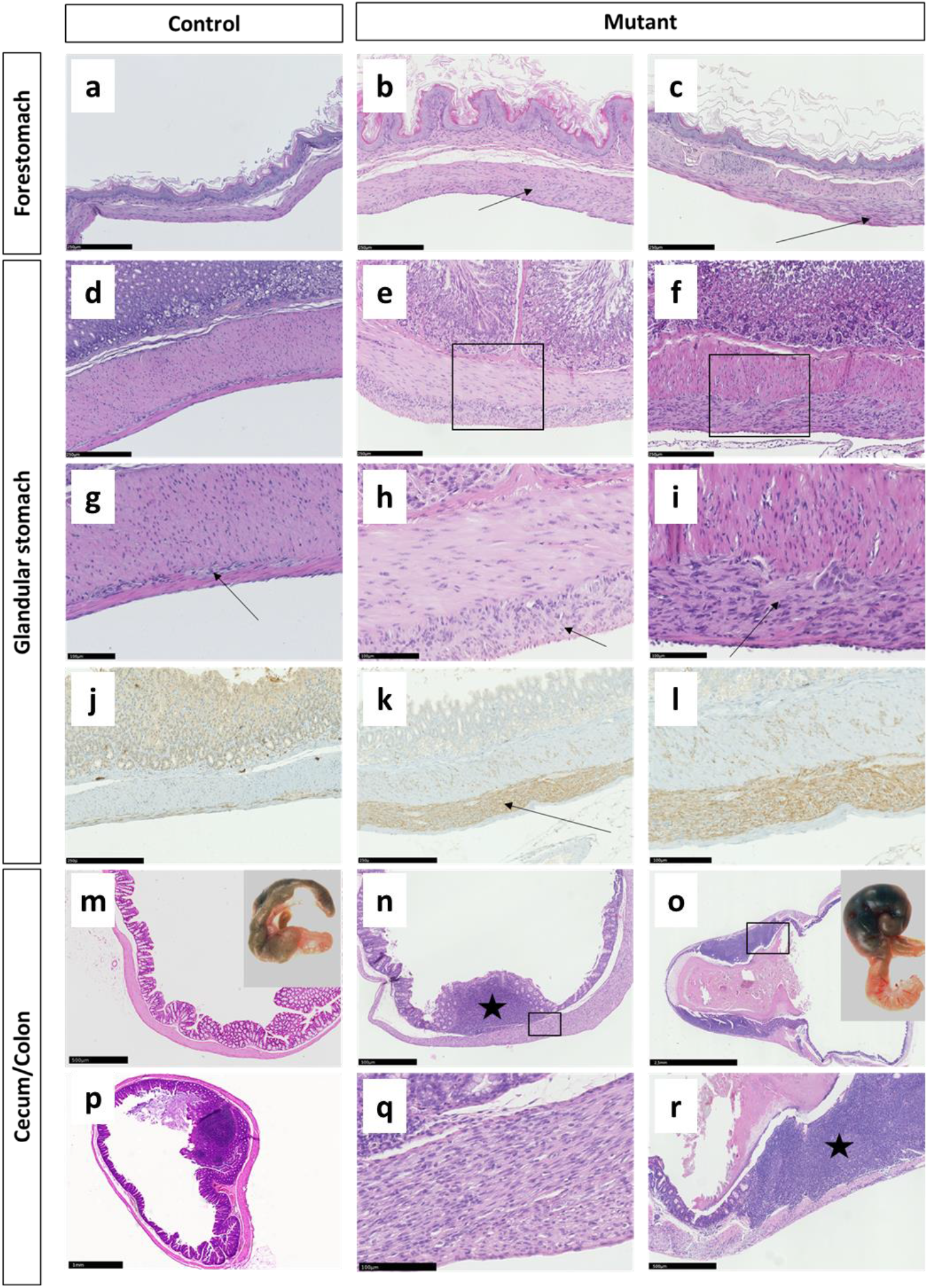
Pathology of the gastrointestinal tract of C3H *Kit*^N824K/WT^ mutant mice: Representative pictures of the forestomach wall of a control (3a) and two mutant animals (3b, 3c) showing a layer of hyperproliferating cells between the inner circular and outer longitudinal muscle layer (arrows), the glandular stomach wall of a control (3d) and two mutant animals (3e, 3f), higher magnification in 3g-3i shows the thin layer of ICCs between the muscular layers (3g, arrow) which is thickened due to ICC hyperplasia in mutant animals (3h, 3i, arrows), IHC for Kit of the gastric wall in a control (3j) and a mutant (3k) animal – higher magnification in 3l - demonstrating Kit-positivity of ICCs in the wild-type and hyperproliferating cells (arrow) in the mutant animal, macroscopic (inserted picture) and histologic appearance of the cecum in a 6 months old wild-type animal (3m) and a 12 months old control (3p) compared to a 6 months old mutant with prominent Peyer’s Patches (star) (3n, 3q) and a 12 months old mutant animal with ingesta retention and extensive inflammation (star) (3o, 3r).

While in three young mutant animals sacrificed at 2 months of age, the cecum was small compared to wild-type animals of the same age (not shown), congestions of ingesta in the cecum was found in almost all heterozygous mutant animals older than 10 months (Fig.3 o). Old animals displayed a megacecum with intense distensions also involving parts of the proximal colon. Histological examination revealed atrophy of the normal mucosal folds and acute inflammatory reactions (Fig. 3 r). Protruding Peyer’s Patches (PPs) were found in the cecum of control animals aged more than five months (Fig. 3p), but were already present in the cecum of mutant animals at 2-5 months of age (Fig. 3 n). In older mutant animals these were flattened due to cecal obstipation (Fig. 3 o, r).

No nodular lesions or tumors comparable to GISTs in humans were detected in C3H *Kit*^N824K/WT^ mice.

### Aged female C3H Kit^N824K/WT^ mutant mice develop mammary carcinomas with high penetrance

With age of 8 months the first subcutaneous tumors were detected in a female *Kit*^N824K/WT^ mutant mouse during necropsy. Out of 24 female mutant animals aged 8 or more months, 18 animals displayed such neoplasms (Fig 4a). Histological analysis revealed these tumors to be Kit-expressing mammary carcinomas, with solid and glandular-like structures (Fig. 4 b, d-e). Transformed mammary epithelial cells were obvious in the subcutaneous areas analyzed, with normal ducts surrounded by lesions corresponding to different mammary transformation stages: hyperplasia, adenoma/mammary intraepithelial neoplasia and early/late carcinoma (Fig. 4 b, c). Immunohistochemical characterization of human breast cancer markers (ER, PR, ErbB2/Her2) in mammary tumors from 14 heterozygous mutant female mice revealed heterogeneity in their expression (Fig. 4d). After Allred scoring, estrogen receptor (ER) was found positive (and mainly cytoplasmic) in 8 out of 14 analyzed tumors (Fig. 4 f,g). Progesterone receptor (PR) was predominantly negative (4h) except for scattered nuclear staining in 4 out of 14 tumors (Fig. 4 i). Herceptest scoring for ErbB2/Her2 expression revealed no positivity as the characteristic ErbB2 membrane staining was mainly detected in stromal (data not shown) but not in tumor cells (Fig. 4 j). Numerous mast cells were observed in the stroma and in the tumor microenvironment (4 k-l). In six out of 18 mutants with mammary tumors, metastases were found in the lung after analyses of H&E-stained sections (Fig. 4 m).

**Figure 4.**
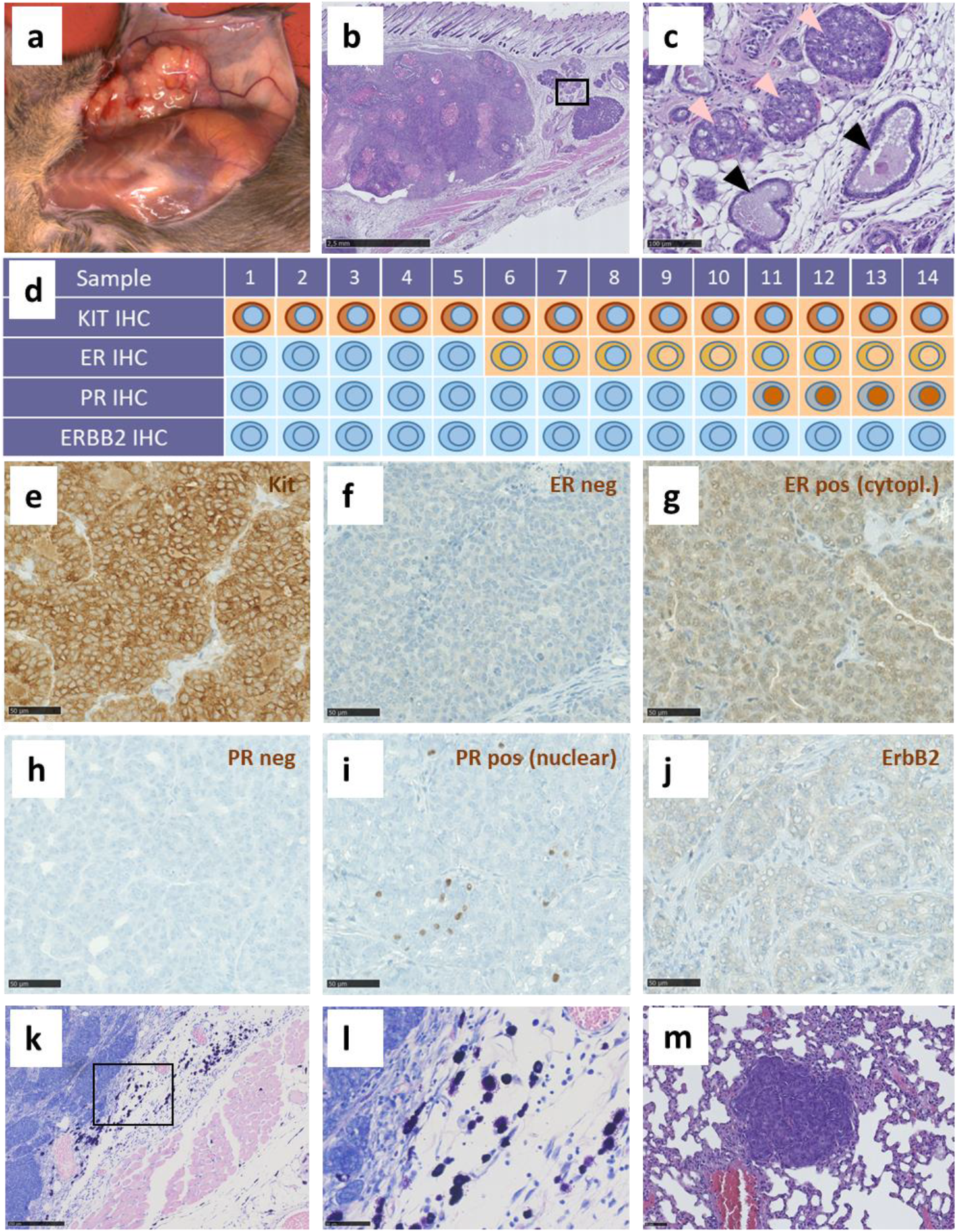
Pathology of mammary tumors found in older female C3H *Kit*^N824K/WT^ mutant mice: Macroscopic (4a) and histological (4b, 4c) appearance of mammary tumors found in older female mutant mice. A high magnification of a region in the periphery of the tumor (4c) show normal mammary ducts (black arrows) and mammary intraepithelial neoplasia (MIN) lesions (bright pink arrows). Overview of immunohistochemistry staining results (negative=blue, positive=brown) and schematic depiction of observed staining patterns (membranous, cytoplasmic, nuclear) and staining intensities (dark brown=strong, light brown=weak) in 14 investigated tumors (4d). Representative IHC pictures of Kit (4e), ER (4f, g), PR (4h, i) and ErbB2 (4j) in the mammary tumors. Examples of a negative (left) and a positive (right) staining for ER and PR are shown. Nuclear ER was also detected in some tumors (4d). Mammary tumor Giemsa staining representative images for the identification of mast cells (4k, l). H&E-stained representative picture of a lung metastasis found in an old mutant female with a mammary tumor (4m).

Further we observed indications of leukosis in one female mutant animal. Liver tumors were found in single male animals from both groups, mutant and control mice. We did not detect melanomas or tumors of the ovaries or testes in any of the mutant animals, but cystic masses at the ovaries of three female control animals. An overview of the macroscopic and histological findings is given in supplementary table 1.

### Macroscopic pathology of four mutant animals and histopathology of one tumor reveal variability of gut pathology with malignant GIST development in mutant N2-hybrid mice

Dissection of four N2-hybrid littermates showing polycythemia from the backcross to C57BL/6JIco at the age of eight months revealed considerable variability concerning the cecum phenotype (Fig. 5 a-d). In one animal, the cecum was similarly small as it was observed in young mutant animals on C3H background, but with a general thickening of the cecal wall and small tumors at the cecum-colon junction (Fig. 5a). One animal showed obstipation of the cecum and mild thickening of the cecal wall, similar to older mutants on C3H background (5b). In the remaining two animals exophytic tumors were found at the cecum (5 c, d). The bigger one was subjected to histopathology, which revealed a malignant GIST phenotype with numerous mitotic figures (Fig 5 e, f).

**Figure 5.**
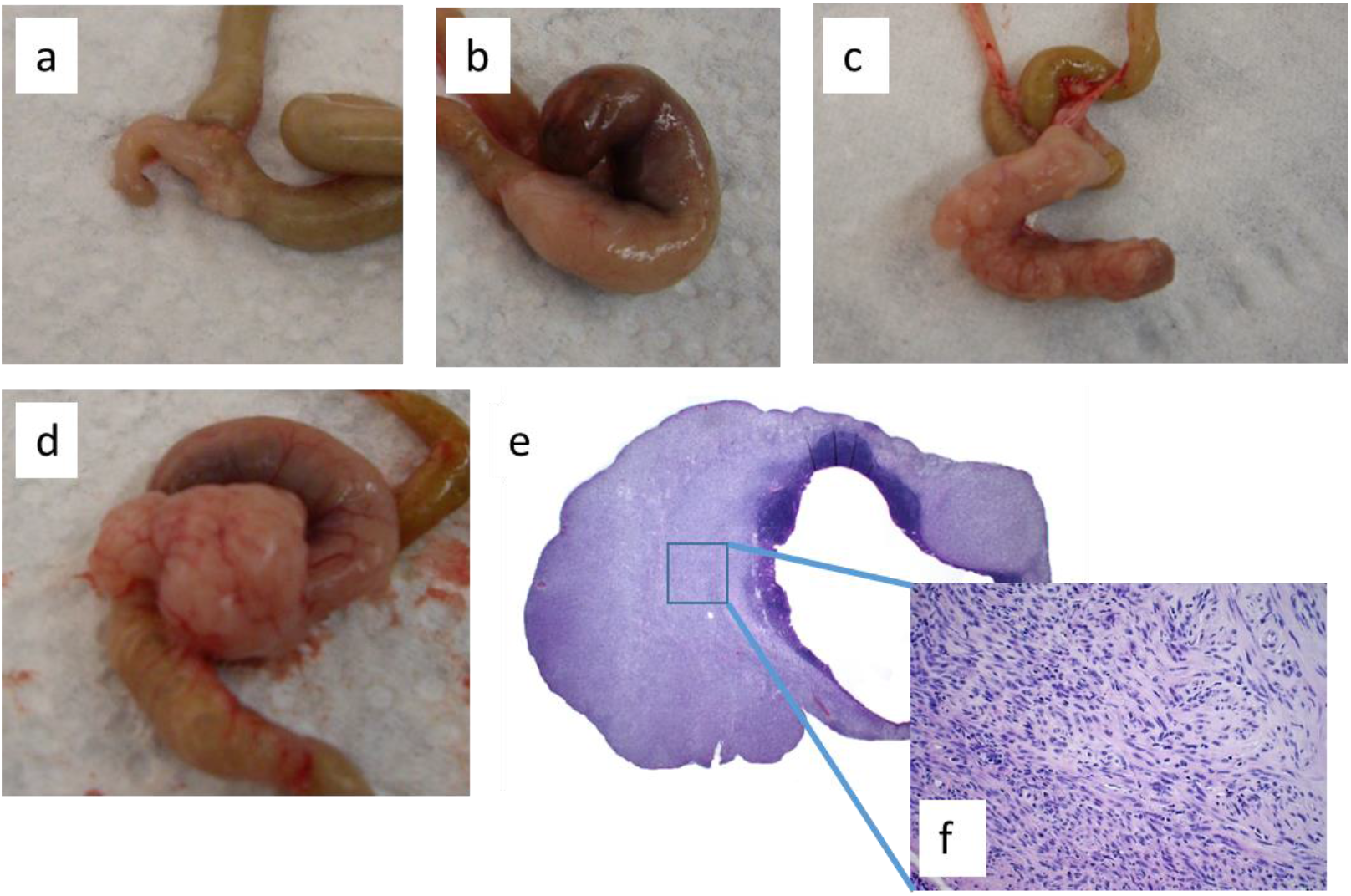
Phenotypic variability of the cecum in mutant N2-hybrid animals: Macroscopic pictures of the cecum derived from four 8 months old N2-hybrid littermates showing polycythemia (5a-5d), histopathology of the tumor depicted in 5d in 10-fold (5e) and 40-fold (5f) magnification.

## Discussion

Gain-of-function mutations in KIT have been found in germ cell tumors, mastocytosis, AML and GIST in humans ^18–21^. Patients with hematopoietic diseases like AML mostly show mutations in the A-loop region of the *KIT* gene, the most prominent ones being mutations at D816 and N822 ^22^. The C3H *Kit*^N824K/WT^ mice carry a *Kit* mutation in the A-loop region, which corresponds to the human N822K mutation first described in an AML cell line with an 8;21 chromosome translocation, named Kasumi 1 ^23^.

The most prominent phenotype found is the microcytic polycythemia with significantly increased hemoglobin and hematocrit values observed in all heterozygous mice. This was associated with increased proliferation rates of bone marrow stem cells in six months old and reduced proliferation in one year old mutant animals. KIT signaling is known to be important for survival and proliferation of erythroid progenitors and so this phenotype is in line with the importance of the A-loop region for hematopoiesis ^6^. Nevertheless, no clear signs of leukemia or increased incidence of leukemic neoplasias as compared to littermate controls were seen. The mutant phenotype rather resembles polycythemia vera (PV) in humans. Interestingly, rare cases of PV patients carrying KIT mutations have been reported ^24^. Additionally, in C3H *Kit*^N824K/WT^ mice an increased proportion of granulocytes in peripheral blood leukocytes and in the spleens of mutant mice was observed. Comparable phenotypes were described for *Kit*^V558Δ;T669I/+^ mice, bearing a mutation in the juxtamembrane domain of KIT and an additional “gatekeeper” second site mutation ^12^. Interestingly, both phenotypes, microcytic erythrocytosis and an increase in granulocytes and monocyte-macrophage populations in the spleen and bone marrow were found in *Kit*^V558Δ;T669I/+^ mice, suggesting that proliferation of the myeloerythroid lineage is affected by this double mutation in a similar way as in the *Kit*^N824K/WT^ single A-loop mutation ^25^.

Further observed in double mutant *Kit*^V558Δ;T669I/+^ mice was a hyperplasia of ICCs in the large intestine and stomach. KIT-activating mutations are found in 60-90% of GIST patients ^26^. Most often these mutations are localized in the juxtamembrane domain of the receptor. Only around 2% of activating point mutations in GISTs reside in the TKII domain in exon 17, where the N822K mutation is located ^5,27^. Most of the mouse models established so far, are characterized by development of GIST lesions at the cecum in heterozygous or homozygous mutant animals ^12–15^. ICC hyperplasia is a generic term of microscopic KIT expressing spindle cell lesions detected during surgery of patients suffering from larger GISTs or other non-GIST lesions ^28,29^. It is described as a thin layer of spindle-shaped proliferating cells between the inner circular and the outer longitudinal muscle layer ^28,30^. Agaimy et al. reported on cases of diffuse Cajal cell hyperplasia with involvement of the complete layer of the muscularis propria in the distal colon, which is an uncommon region for GISTs ^28^. *Kit*^V558Δ/+^ and Kit-K641E mice also develop ICC hyperplasia and neoplastic lesions in the cecum, with *Kit*^V558Δ/+^ mice showing normal cell counts in blood recapitulating the main characteristics of human GIST patients ^14,15^. The C3H *Kit*^N824K/WT^ mouse line mimics a mixture of observations made in humans. The ICC hyperplasia is of 100 % penetrance with stomach, cecum and colon being affected. It is localized either between the two muscular layers or replacing the complete *muscularis propria*. While in humans and also in the double mutant *Kit*^V558Δ;T669I/+^ mice GISTs develop from ICC hyperplasia, there was no tumor progression in the GI tract of Munich C3H *Kit*^N824K/WT^ mice, even in advanced age.

However, *Kit*^N824K/WT^ animals suffer from slowly progressing constipation of the cecum with a penetrance of 100% in animals older than 10 months. ICCs are known to be pacemakers for different motor activities in the gut. In patients suffering from chronic idiopathic constipation or megacolon the number of ICCs is decreased ^31^. In the Munich *Kit*^N824K/WT^ mice the network seems to be disturbed leading to motility problems, although the animals show an ICC hyperplasia. The *Kit*^V558Δ^/+ mice also argue for this because distended ileum and cecum were found in this mutant as well ^15^.

An interesting phenotype we observe in our mice, which to our knowledge has not been described for any other KIT-activating mutation model, is the development of mammary tumors in female mice aged 8 months and older. Since the background strain C3HeB/FeJ does not carry the mammary tumor virus (https://www.jax.org/strain/000658) and age matched control mice display no tumors this pathology is genotype-related. The tumor-type pattern was ranging from solid, sometimes with comedonecrosis, to glandular with partially large lumina. A study of Mayr et al. in 2019 investigated the correlation between GIST and additional neoplasms, and 37% of patients with GIST showed at least one additional tumor, with 20% of these being neoplasms of breast/female genital system. The reason for this association remains unknown ^32^. KIT is highly expressed in normal mammary gland tissue and is described as a progenitor marker in the mouse mammary gland ^33^. However, carcinogenesis is mostly associated with a loss of its expression ^34^. In human breast cancer KIT expression is downregulated during epithelial transformation by KIT promoter hypermethylation ^35^. Other published data suggest that KIT has a dual role in cancer, acting both as an oncogene via its kinase domain or as a tumor suppressor through induction of apoptosis ^36^. Meanwhile an association of strong KIT expression with high-grade breast carcinomas of the basal-like subtype has been shown ^37,38^. However, although a majority of triple-negative mammary carcinomas shows strong expression of KIT, activating mutations of KIT in this type of cancers seem to be rare in humans ^10,39,40^. Our immunohistochemical characterization results of the mammary tumors of this mouse model go in line with those of Eroglu et al. in human breast cancer, where KIT expression was linked to ER positivity but was found independent of ErbB2 ^41^. Our results indicate that activating *Kit* mutations are a potential primary cause of mamma carcinoma development in mice.

While mutations in the A-loop region of KIT are known to be associated to human mastocytosis ^6^, we did not observe an increase in mast cells neither in skin nor in any organ except in the subcutaneous fat surrounding the mammary tumors in female mutant mice. Mast cell (MC) infiltration is known in a range of tumors including breast cancer where they are common components of the tumor stroma. Huang et al. showed that MC infiltration and activation in breast tumors is mainly mediated by stem cell factor and the KIT receptor ^42^. However, the role of increased MCs in cancer is controversial and high numbers of MCs have been associated with good prognosis as well as with poor prognosis and distant metastasis in breast cancer ^43^. Our mice are a valuable model to study the impact of *Kit* mutation on MC infiltration and tumor progression in mammary tumors.

The homologous amino acid exchange at position 822 in humans was also been found in testicular seminoma in humans ^27^. Although we did not observe germ cell tumors in C3H *Kit*^N824K/WT^ mice, the observations of apparently increased fertility of heterozygous mutant animals and the hint towards homozygous lethality could be interesting in this respect. Further studies are required to investigate effects on germ cell development and embryogenesis in detail.

Our model shares several phenotypes described for other rodent *Kit*-mutant models, carrying activating mutations. However, slowly progressing constipation of the cecum and especially development of mammary tumors were not reported for any of these models. Instead, all of the models developing ICC hyperplasia also suffered from GIST tumors at the cecum with ileo-cecal obstruction leading to early killing or spontaneous death. Therefore, mutant animals of these models might not get old enough to develop mammary carcinomas, since these occurred in our model in 8 months old or even older females.

Preliminary results from mice with mixed genetic background suggest that the genetic background significantly affects disease phenotype progression. The *Kit*^N824K/WT^ mice were bred on a pure C3HeB/FeJ background, except for the hybrid animals produced for linkage analysis. While no GIST tumors were found in mutants on C3H background, three out of four mutant hybrid animals sacrificed with an age of eight months showed tumors at the cecum. One of these tumors was histopathologically analyzed and displayed typical signs of malignancy representing GIST. These differences likely result from background-specific variants in modifying genes that influence disease progression in *Kit*^N824K/WT^ mutant mice and could be identified using this model.

The present study has limitations. Since the majority of analyses performed for phenotypic characterization was conducted before the causative mutation was identified, animals for these studies were assigned to the mutant and wild-type groups according to their hematopoietic abnormalities. Since tissue samples were not available later from all of these animals, correct classification could not be confirmed for each individual animal by genotyping. However, genotype assessment of a part of the animals used in the phenotyping studies and of additional animals from the maintenance breeding colony indicated an almost complete match of hematological abnormalities and heterozgous genotype of the causative mutation. Single mismatches could be due to a mix-up of individual mice during sample collection. Further, only a small number of mutant mice younger than five months was examined in pathology, and the same holds true for hybrid mutant animals. Therefore it is difficult to final conclude, whether the abnormally small cecum found in young mutants is a general feature of this model. However, we were able to confirm this observation in the follow-up disease monitoring study (Kraiger et al.). Similarly, we cannot draw final conclusions concerning the effects of genetic background on tumor development in our model from only four mutant animals investigated in this study. Nevertheless, the observation of GIST tumor development in three out of these four mice on the background of numerous investigated mutants on C3H-background not showing this phenotype, clearly indicates a significant influence, that is worth to follow up in future studies. Using this model it could be possible to identify modifiying genes, with protective or oncogenic alleles in different mouse strains.

In summary, the slow, predictable and traceable age-related disease progression of the presented mice bearing an activating Kit mutation render C3H *Kit*^N824K/WT^ mice an essential platform for studying disease modifying genes and assessing novel therapeutic strategies against cancer. It is especially expected to help shedding light on the yet still unclear role of KIT mutations and KIT signaling in mammary carcinoma development.

## Materials and methods

### Mice

MVD013 mice were bred on a pure C3HeB/FeJ (C3H) inbred background by mating heterozygous phenotypic mutants to C3H wild-type (C3H-WT) animals for more than 10 generations before in depth genetic and phenotypic characterization was started. The animals were maintained in the German Mouse Clinic (GMC) in groups of 2-5 animals in individually ventilated cages with standard chow (Breeding diet for mice 1314, Altromin, Lage, Germany) and water accessible *ad libitum* under standard housing conditions (room temperature 23±1 °C, humidity 50±10%) in a 12/12-hour light-dark regime.

Homozygous viability was investigated in the offspring of 20 matings of heterozygous male and female mice. Fertility assessment was done by retrospective analysis of the breeding outcome of maintenance breeding for mouse line MVD013 in the GMC in comparison to maintenance and cohort breeding of the C3HeB/FeJ wild-type colony.

In total, five cohorts of heterozygous mutant mice and wild-type littermates were used for the genetic and phenotypic characterization of this mutant line: One cohort of 100 mice on N2-hybrid mixed genetic background for linkage analysis to map the causative mutation, and 4 cohorts on C3H background for the phenotypic characterization: two cohorts of 8-11 mutant and wild-type animals each per sex were used for the two pipeline phenotyping procedure in the German Mouse Clinic (GMC) ^44^, one cohort of 10 animals of each sex and genotype for the pathology of 12-14 months old animals, and one cohort of 10 six months old female and four 12 months old female animals of each genotype for bone marrow analyses.

Most of the animals used for clinical phenotyping and additional animals from the maintenance breeding colony, from the accompanying disease monitoring study (Kraiger et al., in preparation) and few animals bred for linkage analysis of the mutation were used for macroscopic and histopathological investigations.

All experimental procedures were conducted according to the German Animal Welfare Act, they were approved by the respective authorities of the Government of Upper Bavaria “Regierung von Oberbayern, Sachgebiet 54 – Tierschutz, München, Germany” (Ref. No. 55.2-1-54-2531-78-06; 55.2-1-54-2531-70-07) and carried out and reported in compliance with ARRIVE guidelines.

### Genetic characterization

#### Chromosomal mapping of the mutation by linkage analysis

Heterozygous mutant animals were mated to C57BL/6JIco wild-type mice to generate F1 hybrid animals. Offspring showing the characteristic mutant phenotype of the peripheral complete blood cell count (CBC) were backcrossed to C57BL/6JIco wild-type animals to generate N2 hybrid offspring used for a classical linkage analysis. Animals were grouped according to CBC characteristics in mice showing the mutant, the wild-type or an ambiguous phenotype. Subsequently mice were sacrificed, and DNA isolated from tail tip tissue was used for genome wide SNP genotyping as previously described ^45,46^ followed by classical linkage analysis.

#### Exome sequencing

We performed in-solution targeted enrichment of exonic sequences from spleens of two heterozygous MVD013 mice on C3H background together with six mutants and two wild-type controls from three other lines on the same genetic background using the SureSelectXT Mouse All Exon 50 Mb kit (Agilent Technologies, Santa Clara, CA). The generated libraries were indexed, pooled and sequenced as 100 bp paired-end runs on a HiSeq2000 system (Illumina, San Diego, CA), as previously described ^47^. Read alignment to the mouse genome assembly mm9 was done with Burrows-Wheeler Aligner (BWA, version 0.6.1). Single nucleotide variants (SNV) and small insertions and deletions (indel) were detected with SAMtools (version 0.1.18). Subsequently results were filtered for heterozygous variants present in both MVD013 mutant mice but not in any of the other mice on C3H background analysed in parallel or any of 145 control mice with unrelated phenotypes in the in-house exome sequence database.

#### Genotyping of the mutants

The causative mutation found by exome sequencing in the chromosomal region linked to the mutant phenotype was subsequently confirmed by sequence analysis in two mutant animals and PCR-RFLP based genotyping of heterozygous mutant and wild-type littermates. PCR-RFLP was conducted using 5’-AAACGGGAATATCACTTGCACC-3’ and 5’-CATGTGACATTACAAGGTAGGAG-3’ as primers and the restriction enzyme “Tasl” with the cutting sequence “AATT”. Due to the mutation, the cutting sequence “AATT” in wild-type is changed to “AAAT” in mutants, resulting in differentially sized restriction fragments of 251 nt and 88 nt for the wild-type and 335 nt for the mutant allele. In addition, genotyping by heteroduplex detection was performed with a LightScanner^®^ device originally from Idaho Technology Inc. (distributed by Bioke, Leiden, Netherlands) as described previously ^48^, from PCR-products of the respective region obtained with the following primer pair: forward TTCCTGTGAATGGAAGGAAG, and reverse AAAGCACCCTGGGTAGACTC.

#### Systematic phenotyping at the GMC

A systematic GMC phenotype screen using a two-pipeline procedure as described by Fuchs et al. 2012 and additional pathological investigations on aged animals were carried out for clinical phenotype assessment of mutant mice ^44^. Animals were derived from heterozygous mutant to wild-type matings and assigned as polycythemic mutant and normal controls according to their hematological phenotype determined at 10-12 weeks, since the causative mutation was still unknown. Groups of eight to eleven male and female mice of each phenotype (mutant or wild-type) were analyzed in each test, starting with first tests at age 14-15 weeks and finishing at 22-23 weeks. Only tests which detected polycythemia-associated phenotypes are described in detail.

#### Blood collection

Blood collection: Blood samples obtained by puncture of the retrobulbar vein plexus from mice anesthetized with isoflurane were collected in Li-heparin coated tubes (Kabe Labortechnik GmbH, Nümbrecht-Elsenroth, Germany) for clinical chemistry analyses and EDTA-coated tubes (Kabe Labortechnik GmbH, Nümbrecht-Elsenroth, Germany) for the determination of hematological values. Heparinized blood samples were stored at room temperature for 1-2 hours before being separated by centrifugation (5000 x g, 10 minutes, 8°C) and plasma transferred to plain sample tubes for analysis.

#### Clinical chemistry analyses

Plasma samples were used diluted 1:2 with deionized water to determine plasma parameters with an AU400 analyzer (Olympus Deutschland GmbH, Hamburg, Germany) as described ^49^.

#### Hematological investigations

Hematological investigations: EDTA-blood samples were agitated at room temperature on a rotary until analyzed with an abc-animal blood counter (Scil Animal Care Company, Viernheim, Germany) to determine parameters of the peripheral complete blood count (CBC) including automated cell size-dependent pre-differentiation of leukocytes.

#### FACS analysis of peripheral blood leukocytes

FACS analysis of peripheral blood leukocytes: Subpopulations of peripheral blood leukocytes were analyzed using a BD LSR II flow cytometer (BD Biosciences, San Jose, USA) and fluorescence-conjugated antibodies against CD45, CD3, CD4, CD8, CD19, CD25, B220, CD5, IgD, CD11b, Gr-1, MCH-class II, Ly6C, CD44 and CD62L as described previously ^50^.

### Analysis of bone marrow characteristics

#### FACS analysis of hematopoietic stem and progenitor cells

Total bone marrow (BM) from one femur and two tibiae was isolated by flushing. After cell counting 1×10^6^ cells were stained with a cocktail for lineage markers (CD3, CD11b, B220, Ter119, and Ly6G/6C; BioLegend), and lineage-negative cells (Lin-) were analyzed for c-Kit (clone 2B8; eBioscience) and Sca-1 (clone D7; BioLegend) expression to discriminate Kit+ Sca-1- (KL) and Kit+ Sca-1+ (KSL) cells. Progenitor and stem cell subpopulations were identified by staining with the following additional markers: CD34 (clone MEC14.7; BioLegend), Fc-gamma-II/III-R (clone 93; eBioscience).

#### Histology of bone marrow

Femurs were fixed in 10% formalin, subsequently decalcified in 10% buffered EDTA, pH 7.2, and afterwards embedded in paraffin. Sections were stained with hematoxylin and eosin (H&E) for histological analysis.

#### Additional phenotyping methods

Further analyses during systemic phenotype assessment are described in supplement 1, and additional analyses on proliferation of bone marrow cells and apoptosis rates of stem cells can be found in supplement 2.

#### Statistical analyses

Data displayed in Fig. 1 and Fig. S1 were evaluated separately for males and females by a Wilcoxon-Rank-Sum test performed with GraphPad Prism. In general, data obtained from systematic phenotype analyses (Supplement 1 and tables therein) were analyzed by T-test or Wilcoxon-Rank-Sum-Test for genotype-effects and, if appropriate, by 2-way ANOVA analysis for effects of sex and genotype. The level of significance was set at p<0.05. We do not apply multiple testing correction during evaluation of primary phenotyping results, since this analysis is a screening procedure for the generation of hypotheses.

#### Macroscopic pathology, histology and immunohistochemistry

In total we investigated 106 animals on C3H background macroscopically (76 heterozygous mutant and 30 wild-type littermates) and mutants from one litter N2-hybrid animals (n=4). Organs of 60 mutant animals on C3H background and the cecum of one N2-hybrid mutant were subjected to histopathological analysis. During necropsy macroscopical analysis was performed previous to histological examination of H&E stained, formalin-fixed, paraffin-embedded tissues. Images were taken by the slide scanning system NanoZoomer^®^ 2.O HT (Hamamatsu, Japan). Giemsa staining was carried out for visualization of mastocytes in 3 μm paraffin sections of 8 mutant and 4 control animals. All organs were analysed, including tumors of female mice.

Immunohistochemical staining was carried out on 1 μm tissue section in automated immunostainers (Discovery®XT, Roche, Penzberg, Germany, and Leica Bond, Wetzlar, Germany) using the streptavidin-peroxidase method. The following primary antibodies were used: anti-B220 (catalog no.550286, 1:50, BD PharMingen, California, USA), anti-KIT antibody (catalog no. AF1356, 1:40, R&D Systems,Minnesota, USA), anti-CD3 (catalog no. ZYT-RBG024, 1:8, diagnostic-Biosystems, California, USA), anti-estrogen receptor alpha (catalog no. ab75635, 1:100, Abcam, Berlin, Germany) anti-ErbB2 (catalog no. RB-103-P0200 1:200, Thermo Scientific, Massachusetts, USA), anti-myeloperoxidase (MPO) (catalog no. ab208670, 1:1200, Abcam, Berlin, Germany), anti-progesterone receptor (catalog no. RM-9102-S0, 1:400, Thermo Scientific) and anti-TER-119 (catalog no. 550565, 1:50, BD PharMingen, California, USA). To confirm antibody specificity positive controls with known protein expression as well as negative controls without primary antibody were used. The slides were analyzed independently by two pathologists.

## Supporting information

Kit-N824K mutant Supplemental data

## German Mouse Clinic Consortium

Juan Antonio Aguilar Pimentel^1^

Lore Becker^1^

Lillian Garrett^1,7^

Sabine M. Hölter^1,7,8^

Cornelia Prehn^1,9^

Ildikó Rácz^1,10++^

Jan Rozman^1##^

Oliver Puk^1,7xx^

Anja Schrewe^1^

Holger Schulz^11^

Jerzy Adamski^1,12,13^

Dirk H. Busch^14^

Irene Esposito^15§§^

Wolfgang Wurst^7,8,16,17^

Claudia Stoeger^1^

^1^Institute of Experimental Genetics, German Mouse Clinic, Helmholtz Zentrum München, German Research Center for Environmental Health, Neuherberg, Germany

^7^Institute of Developmental Genetics, Helmholtz Zentrum München, German Research Center for Environmental Health, Neuherberg, Germany

^8^Chair of Developmental Genetics, TUM School of Life Sciences, Technische Universität München, Freising-Weihenstephan, Germany

^9^Metabolomics and Proteomics Core, Helmholtz Zentrum München, German Research Center for Environmental Health, Neuherberg, Germany

^10^Clinic for Neurodegenerative Diseases and Geriatric Psychiatry, Medical Faculty, University of Bonn, Bonn, Germany

^11^Institute of Lung Biology and Disease, Helmholtz Zentrum München, German Research Center for Environmental Health, Neuherberg, Germany

^12^Department of Biochemistry, Yong Loo Lin School of Medicine, National University of Singapore, Singapore, Singapore

^13^Institute of Biochemistry, Faculty of Medicine, University of Ljubljana, Ljubljana, Slovenia

^14^Institute for Medical Microbiology, Immunology and Hygiene, Technische Universität München, Munich, Germany

^15^Institute of Pathology, Helmholtz Zentrum München, German Research Center for Environmental Health, Neuherberg, Germany

^16^Deutsches Institut für Neurodegenerative Erkrankungen (DZNE) Site Munich, Munich, Germany

^17^Munich Cluster for Systems Neurology (SyNergy), Ludwig-Maximilians-Universität München, Munich, Germany

^++^Current address: Clinic of Neurodegenerative Diseases and Gerontopsychiatry, University of Bonn Medical Center, Bonn, Germany

^##^Current address: Institute of Molecular Genetics of the Czech Academy of Sciences, Czech Centre for Phenogenomics, Vestec, Czech Republic

^xx^Current address: Praxis für Humangenetik Tübingen, Tübingen, Germany

^§§^Current address: Institute of Pathology, Universitätsklinikum Düsseldorf, Düsseldorf, Germany

## Acknowledgments

We acknowledge the excellent support of the technicians and animal caretakers of the German Mouse Clinic. Special thanks to the technicians of the pathology lab.

## Author Contributions

B.R., K.M., J.C.-W., V. G.-D., H.F., H.P. were involved in study design, S.S. and B.R. established the mouse model, T. K.-R., A.G. and B.R. drafted the manuscript and contributed and analysed data, K.M., A S.-M., T.A., M.Kl., B.A., A.G., H.P., J.A.A.P., L.B., L.G., S.M.H., C.P., I.R., J.R., O.P., A.S., H.S., J.A., D.H.B., I.E., W.W. contributed data and were involved in data analyses and interpretation, A. S.-M., B.A., M.Kr., V.G.-D., E.W., A.G., M. H.dA. and C.S. contributed to data interpretation and writing of the manuscript, E.W., W.W. and M.HdA. provided funding and infrastructure. All authors contributed to the article and read and approved the submitted version.

## Data availability statement

All phenotypic data generated and analyzed during the current study are included as graph and/or tables in this published article and its supplementary information files. Result overviews of genetic analyses (linkage analysis, Exome sequencing) are also displayed in the supplementary data file. Raw data of SNP genotyping-based linkage analysis, exome sequencing results and additional histopathology images are stored in in-house repositories at the Helmholtz Center Munich and will be made available from the co-author H. Fuchs on reasonable request.

## References

1 Russell, E. S. Hereditary anemias of the mouse: a review for geneticists. Advances in genetics 20, 357–459 (1979).

2 Kitamura, Y. & Go, S. Decreased production of mast cells in S1/S1d anemic mice. Blood 53, 492–497 (1979).

3 Huizinga, J. D. et al. W/kit gene required for interstitial cells of Cajal and for intestinal pacemaker activity. Nature 373, 347–349, doi:10.1038/373347a0 (1995).

4 Min, K. W. & Leabu, M. Interstitial cells of Cajal (ICC) and gastrointestinal stromal tumor (GIST): facts, speculations, and myths. Journal of cellular and molecular medicine 10, 995–1013 (2006).

5 Rubin, B. P. et al. KIT activation is a ubiquitous feature of gastrointestinal stromal tumors. Cancer Res 61, 8118–8121 (2001).

6 Boissan, M., Feger, F., Guillosson, J. J. & Arock, M. c-Kit and c-kit mutations in mastocytosis and other hematological diseases. J Leukoc Biol 67, 135–148 (2000).

7 Lennartsson, J., Jelacic, T., Linnekin, D. & Shivakrupa, R. Normal and oncogenic forms of the receptor tyrosine kinase kit. Stem Cells 23, 16–43, doi:10.1634/stemcells.2004-0117 (2005).

8 Kemmer, K. et al. KIT mutations are common in testicular seminomas. Am J Pathol 164, 305–313, doi:10.1016/S0002-9440(10)63120-3 (2004).

9 Kanapathy Pillai, S. K., Tay, A., Nair, S. & Leong, C. O. Triple-negative breast cancer is associated with EGFR, CK5/6 and c-KIT expression in Malaysian women. BMC Clin Pathol 12, 18, doi:10.1186/1472-6890-12-18 (2012).

10 Simon, R. et al. KIT (CD117)-positive breast cancers are infrequent and lack KIT gene mutations. Clin Cancer Res 10, 178–183, doi:10.1158/1078-0432.ccr-0597-3 (2004).

11 Spitaleri, G. et al. Inactivity of imatinib in gastrointestinal stromal tumors (GISTs) harboring a KIT activation-loop domain mutation (exon 17 mutation pN822K). Onco Targets Ther 8, 1997–2003, doi:10.2147/OTT.S81558 (2015).

12 Bosbach, B. et al. Imatinib resistance and microcytic erythrocytosis in a KitV558Delta;T669I/+ gatekeeper-mutant mouse model of gastrointestinal stromal tumor. Proc Natl Acad Sci U S A 109, E2276–2283, doi:10.1073/pnas.1115240109 (2012).

13 Nakai, N. et al. A mouse model of a human multiple GIST family with KIT-Asp820Tyr mutation generated by a knock-in strategy. The Journal of pathology 214, 302–311, doi:10.1002/path.2296 (2008).

14 Rubin, B. P. et al. A knock-in mouse model of gastrointestinal stromal tumor harboring kit K641E. Cancer research 65, 6631–6639, doi:10.1158/0008-5472.CAN-05-0891 (2005).

15 Sommer, G. et al. Gastrointestinal stromal tumors in a mouse model by targeted mutation of the Kit receptor tyrosine kinase. Proc Natl Acad Sci U S A 100, 6706–6711, doi:10.1073/pnas.1037763100 (2003).

16 Aigner, B. et al. Generation of N-ethyl-N-nitrosourea-induced mouse mutants with deviations in hematological parameters. Mamm Genome 22, 495–505, doi:DOI 10.1007/s00335-011-9328-4 (2011).

17 Lennartsson, J. & Ronnstrand, L. Stem cell factor receptor/c-Kit: from basic science to clinical implications. Physiol Rev 92, 1619–1649, doi:10.1152/physrev.00046.2011 (2012).

18 Shimada, A. et al. N822 mutation of KIT gene was frequent in pediatric acute myeloid leukemia patients with t(8;21) in Japan: a study of the Japanese childhood AML cooperative study group. Leukemia 21, 2218–2219, doi:10.1038/sj.leu.2404766 (2007).

19 Hersmus, R. et al. Prevalence of c-KIT mutations in gonadoblastoma and dysgerminomas of patients with disorders of sex development (DSD) and ovarian dysgerminomas. PLoS One 7, e43952, doi:10.1371/journal.pone.0043952 (2012).

20 Baek, J. O. et al. N822K c-kit mutation in CD30-positive cutaneous pleomorphic mastocytosis after germ cell tumour of the ovary. Br J Dermatol 166, 1370–1373, doi:10.1111/j.1365-2133.2012.10816.x (2012).

21 Yang, J. et al. Genetic aberrations of gastrointestinal stromal tumors. Cancer 113, 1532–1543, doi:10.1002/cncr.23778 (2008).

22 Qin, Y. Z. et al. Heterogeneous prognosis among KIT mutation types in adult acute myeloid leukemia patients with t(8;21). Blood Cancer J 8, 76, doi:10.1038/s41408-018-0116-1 (2018).

23 Larizza, L., Magnani, I. & Beghini, A. The Kasumi-1 cell line: a t(8;21)-kit mutant model for acute myeloid leukemia. Leukemia & lymphoma 46, 247–255, doi:10.1080/10428190400007565 (2005).

24 Fontalba, A. et al. Identification of c-Kit gene mutations in patients with polycythemia vera. Leukemia research 30, 1325–1326, doi:10.1016/j.leukres.2005.12.020 (2006).

25 Deshpande, S. et al. KIT receptor gain-of-function in hematopoiesis enhances stem cell self-renewal and promotes progenitor cell expansion. Stem Cells 31, 1683–1695, doi:10.1002/stem.1419 (2013).

26 Longley, B. J., Reguera, M. J. & Ma, Y. Classes of c-KIT activating mutations: proposed mechanisms of action and implications for disease classification and therapy. Leukemia research 25, 571–576 (2001).

27 Kitamura, Y. & Hirotab, S. Kit as a human oncogenic tyrosine kinase. Cellular and molecular life sciences: CMLS 61, 2924–2931, doi:10.1007/s00018-004-4273-y (2004).

28 Agaimy, A. et al. Sporadic segmental Interstitial cell of cajal hyperplasia (microscopic GIST) with unusual diffuse longitudinal growth replacing the muscularis propria: differential diagnosis to hereditary GIST syndromes. Int J Clin Exp Pathol 3, 549–556 (2010).

29 Kindblom, L. G., Remotti, H. E., Aldenborg, F. & Meis-Kindblom, J. M. Gastrointestinal pacemaker cell tumor (GIPACT): gastrointestinal stromal tumors show phenotypic characteristics of the interstitial cells of Cajal. Am J Pathol 152, 1259–1269 (1998).

30 Hirota, S. et al. Cause of familial and multiple gastrointestinal autonomic nerve tumors with hyperplasia of interstitial cells of Cajal is germline mutation of the c-kit gene. The American journal of surgical pathology 24, 326–327 (2000).

31 Wedel, T. et al. Enteric nerves and interstitial cells of Cajal are altered in patients with slow-transit constipation and megacolon. Gastroenterology 123, 1459–1467, doi:10.1053/gast.2002.36600 (2002).

32 Mayr, P. et al. Malignancies associated with GIST: a retrospective study with molecular analysis of KIT and PDGFRA. Langenbecks Arch Surg 404, 605–613, doi:10.1007/s00423-019-01773-2 (2019).

33 Regan, J. L. et al. c-Kit is required for growth and survival of the cells of origin of Brca1-mutation-associated breast cancer. Oncogene 31, 869–883, doi:10.1038/onc.2011.289 (2012).

34 Chui, X. et al. Immunohistochemical expression of the c-kit proto-oncogene product in human malignant and non-malignant breast tissues. British journal of cancer 73, 1233–1236 (1996).

35 Janostiak, R., Vyas, M., Cicek, A. F., Wajapeyee, N. & Harigopal, M. Loss of c-KIT expression in breast cancer correlates with malignant transformation of breast epithelium and is mediated by KIT gene promoter DNA hypermethylation. Exp Mol Pathol 105, 41–49, doi:10.1016/j.yexmp.2018.05.011 (2018).

36 Wang, H. et al. The Proto-oncogene c-Kit Inhibits Tumor Growth by Behaving as a Dependence Receptor. Mol Cell 72, 413–425 e415, doi:10.1016/j.molcel.2018.08.040 (2018).

37 Amin, M. M., El-Hawary, A. K. & Farouk, O. Relation of CD117 immunoreactivity and microvascular density in invasive breast carcinoma. Indian journal of pathology & microbiology 55, 456–460, doi:10.4103/0377-4929.107780 (2012).

38 Tsuda, H. et al. Correlation of KIT and EGFR overexpression with invasive ductal breast carcinoma of the solid-tubular subtype, nuclear grade 3, and mesenchymal or myoepithelial differentiation. Cancer science 96, 48–53, doi:10.1111/j.1349-7006.2005.00009.x (2005).

39 Zhu, Y. et al. C-kit and PDGFRA gene mutations in triple negative breast cancer. International journal of clinical and experimental pathology 7, 4280–4285 (2014).

40 Hussain, S. R. et al. Identification of the c-kit gene mutations in biopsy tissues of mammary gland carcinoma tumor. Journal of the Egyptian National Cancer Institute 24, 97–103, doi:10.1016/j.jnci.2011.10.003 (2012).

41 Eroglu, A. & Sari, A. Expression of c-kit proto-oncogene product in breast cancer tissues. Med Oncol 24, 169–174, doi:10.1007/BF02698036 (2007).

42 Huang, B. et al. SCF-mediated mast cell infiltration and activation exacerbate the inflammation and immunosuppression in tumor microenvironment. Blood 112, 1269–1279, doi:10.1182/blood-2008-03-147033 (2008).

43 Carpenco, E. et al. Mast Cells as an Indicator and Prognostic Marker in Molecular Subtypes of Breast Cancer. In Vivo 33, 743–748, doi:10.21873/invivo.11534 (2019).

44 Fuchs, H. et al. Innovations in phenotyping of mouse models in the German Mouse Clinic. Mamm Genome 23, 611–622, doi:10.1007/s00335-012-9415-1 (2012).

45 Klaften, M. & Hrabe de Angelis, M. ARTS: a web-based tool for the set-up of high-throughput genome-wide mapping panels for the SNP genotyping of mouse mutants. Nucleic Acids Res 33, W496–500, doi:10.1093/nar/gki430 (2005).

46 Herbach, N. et al. Dominant-negative effects of a novel mutated Ins2 allele causes early-onset diabetes and severe beta-cell loss in Munich Ins2C95S mutant mice. Diabetes 56, 1268–1276, doi:10.2337/db06-0658 (2007).

47 Treise, I. et al. Defective immuno- and thymoproteasome assembly causes severe immunodeficiency. Sci Rep 8, 5975, doi:10.1038/s41598-018-24199-0 (2018).

48 Sabrautzki, S. et al. Point mutation of Ffar1 abrogates fatty acid-dependent insulin secretion, but protects against HFD-induced glucose intolerance. Mol Metab 6, 1304–1312, doi:10.1016/j.molmet.2017.07.007 (2017).

49 Stein, C. et al. Clinical chemistry of human FcRn transgenic mice. Mamm Genome 23, 259–269, doi:10.1007/s00335-011-9379-6 (2012).

50 Fuchs, H. et al. Mouse phenotyping. Methods 53, 120–135, doi:10.1016/j.ymeth.2010.08.006 (2011).

