## Supplementary material for "New C3H *Kit*^N824K/WT^ cancer mouse model develops late-onset malignant mammary tumors with high penetrance": Kit-N824K mutant Supplemental data

### Supplement 1: Overview of methods and results from systematic phenotyping in the GMC
### (tests with significant differences between mutants and controls)

#### Bone density analysis

*Equipment:* pDEXA Sabre X-ray Bone Densitometer (Norland Medical Systems. Inc., Basingstoke, Hampshire, UK; distributed by Stratec Medizin-technik GmbH, Pforzheim, Germany). *Quality control***:** Calibration of the system was done in daily intervals using the QC and the QA phantoms delivered by the manufacturer. Results from the quality control were recorded by the system. *Settings:* Scan speed 20 mm/s, Resolution 0.5 mm x 1.0 mm, HAW 0.020. *Procedure:* After anesthesia, the weight and length of the mouse were recorded, and the mouse was placed in the analyzer. After a scout scan, the area of interest was optimized and the measure scan started.

*Data-analysis:* For analysis of the data, regions have to be defined. The standard analysis comprises a whole body analysis as well as a whole body analysis excluding the skull.

**Table 1: Bone- and weight-related quantitative parameters**
(data presented as mean ± standard error of mean)

| **20-week old mice** | **MVD013 control** | | **MVD013 mutant** | | **co ~ mut** | **co ~ mut** | **ANOVA** | | |
|---|---|---|---|---|---|---|---|---|---|
| **Parameter** | **Male** (n=8) | **Female** (n=10) | **Male** (n=10) | **Female** (n=10) | **Male** *p – value* | **Female** *p – value* | *p – value genotype* | *p – value sex* | *p – value interaction* |
| **BMD** $[mg/cm^2]$ | 57 ± 2 | 57 ± 2 | 54 ± 1 | 49↓ ± 1 | n.s. | < 0.001 | < 0.001 | n.s. | n.s. |
| **sBMD** $[10^{-3}$ x $cm^{-2}]$ | 1.79 ± 0.08 | 1.92 ± 0.02 | 1.72 ± 0.04 | 1.84 ± 0.04 | n.a. | n.a. | n.s. | < 0.05 | n.s. |
| **BMC** [mg] | 581 ± 57 | 627 ± 53 | 628 ± 49 | 467 ± 20 | n.a. | n.a. | n.s. | n.s. | < 0.05 |
| **Bone Content** [%] | 1.78 ± 0.13 | 2.09 ± 0.10 | 1.98 ± 0.14 | 1.75 ± 0.06 | n.a. | n.a. | n.s. | n.s. | < 0.05 |
| **Body Length** [cm] | 10.44 ± 0.06 | 10.35 ± 0.08 | 10.50 ± 0.00 | 10.20 ± 0.08 | n.a. | n.a. | n.s. | < 0.01 | n.s. |
| **Body Weight** [g] | 32.28 ± 1.18 | 29.67 ± 1.17 | 31.57 ± 0.62 | 26.63 ± 0.66 | n.a. | n.a. | n.s. | < 0.001 | n.s. |
| **Fat mass** [g] | 5.92 ± 1.39 | 7.12 ± 1.45 | 7.25 ± 0.94 | 3.32 ± 0.57 | n.a. | n.a. | n.s. | n.s. | < 0.05 |
| **Fat Content** [%] | 17.47 ± 3.82 | 22.80 ± 3.92 | 22.80 ± 2.75 | 12.34 ± 1.97 | n.a. | n.a. | n.s. | n.s. | < 0.05 |
| **Lean mass** [g] | 19.20 ± 0.66 | 15.75 ± 0.75 | 17.03 ± 0.93 | 16.85 ± 0.69 | n.a. | n.a. | n.s. | < 0.05 | < 0.05 |
| **Lean Content** [%] | 60.29 ± 3.68 | 54.17 ± 3.68 | 54.05 ± 2.93 | 63.33 ± 2.09 | n.a. | n.a. | n.s. | n.s. | < 0.05 |

#### Intraperitoneal Glucose-Tolerance-Test (age 19 weeks)

The mice were fasted overnight. In the beginning of the test body weight of mice was determined. For the determination of the baseline blood glucose level of the fasted mouse, a small drop of blood collected from the tail vein was analyzed with the Accu-Chek Aviva glucose analyzer (Roche/ Mannheim). Thereafter mice were injected intraperitoneally with 2 g of glucose/kg body weight using a 20% glucose solution, a 25 gauge needle and a 1 ml syringe. 15, 30, 60, 90 and 120 minutes after glucose injection, additional blood samples (one drop each) were collected and used to determine blood glucose levels as described before.

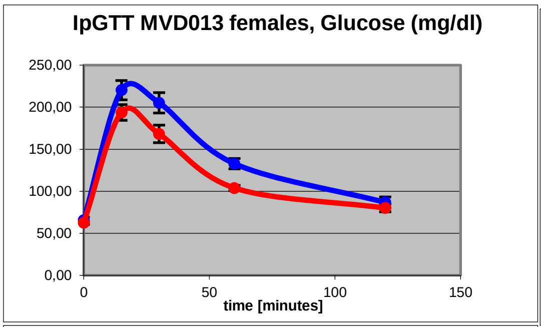


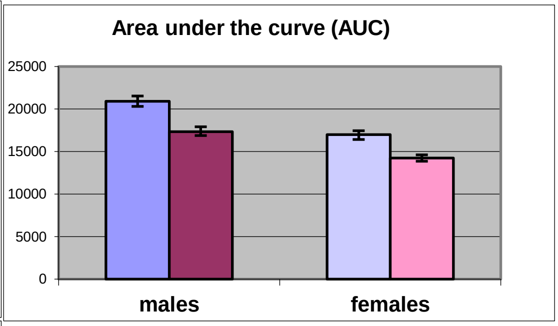


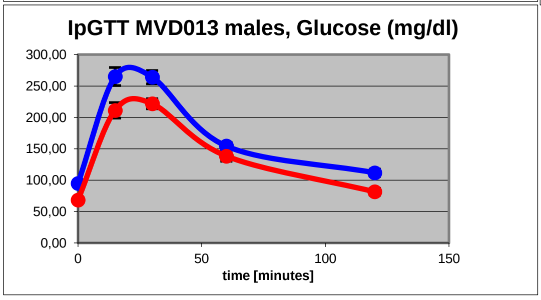


Control: blue, purple; Mutant: red, dark red, pink
AUC significantly decreased in mutants: T-test p<0.001 in males, p<0.01 in females

#### Clinical Chemistry and hematology

Clinical Chemistry analysis of Li-heparin plasma samples. Plasma values of fasted mice (table19) were measured from samples collected after overnight food withdrawal (age 21 weeks). First and second sample (tables 20 -21 and 22-23) were collected from *ad libitum* fed animals at age 20 and 23 weeks.

**Table 19: Blood lipid and glucose values of fasted mice.**
Data are presented as mean ± standard error of mean.

| Parameter | Control (A) | | Mutant (B) | | A~B | A~B | ANOVA | | |
|---|---|---|---|---|---|---|---|---|---|
| | Male | Female | Male | Female | Male | Female | genotype | sex | inter. |
| | (n=8) | (n=10) | (n=10) | (n=10) | *p-value* | *p-value* | *p-value* | *p-value* | *p-value* |
| Cholesterol [mg/dl] | 182.8 ± 2.87 | 129.3 ± 4.04 | 168.9 ± 3.57 | 109 ± 7.98 | p<0.01 | p<0.05 | p<0.01 | p<0.001 | n.s. |
| Triglycerides [mg/dl] | 197 ± 18.5 | 254 ± 21.8 | 166 ± 10.9 | 195 ± 23.9 | n.s. | n.s. | p<0.05 | p<0.05 | n.s. |
| NEFA [mmol/l] | 1.8 ± 0.2 | 2.7 ± 0.13 | 2.1 ± 0.06 | 2.3 ± 0.16 | n.s. | n.s. | n.s. | p<0.001 | p<0.05 |
| Glucose [mg/dl] | 141.6 ± 10.5 | 99.8 ± 8.3 | 104 ± 11.9 | 72.7 ± 6.4 | p<0.05 | p<0.05 | p<0.01 | p<0.001 | n.s. |

| Table 20: Clinical-chemical parameters, 1st sample. Data are presented as mean ± standard error of mean. | | | | | | | | | |
|---|---|---|---|---|---|---|---|---|---|
| Parameter | Control (A) | | Mutant (B) | | A~B | A~B | ANOVA | | |
| | Male | Female | Male | Female | Male | Female | genotype | sex | inter. |
| | (n=11) | (n=9) | (n=9) | (n=11) | *p-value* | *p-value* | *p-value* | *p-value* | *p-value* |
| Sodium [mmol/l] | 154.7 ± 2.06 | 149.6 ± 0.44 | 150.9 ± 0.48 | 148.9 ± 0.49 | n.s. | n.s. | p=0.053 | p<0.01 | n.s. |
| Potassium [mmol/l] | 4.76 ± 0.08 | 3.84 ± 0.08 | 4.75 ± 0.08 | 4.27 ± 0.12 | n.s. | p<0.01 | n.s. | p<0.001 | n.s. |
| Chloride [mmol/l] | 115.8 ± 1.42 | 111.2 ± 0.58 | 114.8 ± 0.48 | 113.9 ± 0.61 | n.s. | p<0.01 | n.s. | p<0.01 | p<0.05 |
| Total Protein [g/dl] | 5.71 ± 0.11 | 5.36 ± 0.06 | 5.49 ± 0.05 | 5.4 ± 0.05 | n.s. | n.s. | n.s. | p<0.01 | p=0.056 |
| Albumin [g/dl] | 2.891 ± 0.062 | 2.911 ± 0.035 | 2.711 ± 0.035 | 2.891 ± 0.031 | p<0.05 | n.s. | p=0.051 | p=0.051 | p=0.051 |
| Creatinine [mg/dl] | 0.08 ± 0 | 0.07 ± 0 | 0.08 ± 0 | 0.08 ± 0 | n.s. | n.s. | p<0.05 | n.s. | n.s. |
| Triglycerides [mg/dl] | 318 ± 23.2 | 255 ± 30.8 | 241 ± 21 | 189 ± 21.5 | p<0.05 | n.s. | p<0.01 | p<0.05 | n.s. |
| NEFA [mmol/l] | 2.1 ± 0.1 | 1.8 ± 0.15 | 2.1 ± 0.08 | 1.7 ± 0.09 | n.s. | n.s. | n.s. | p<0.01 | n.s. |
| ASAT [U/l] | 42.7 ± 3.45 | 55.6 ± 2.53 | 42 ± 1.63 | 72.2 ± 6.38 | n.s. | p<0.05 | n.s. | p<0.001 | p<0.05 |
| Ferritin [ng/ml] | 20.5 ± 1.84 | 26.6 ± 1.57 | 20.2 ± 1.84 | 23.3 ± 2.9 | n.s. | n.s. | n.s. | p=0.053 | n.s. |
| Transferrin [mg/dl] | 164.1 ± 3.44 | 154.2 ± 2.79 | 149.5 ± 1.65 | 147.4 ± 1.03 | p<0.01 | p<0.05 | p<0.001 | p<0.01 | p<0.05 |
| Iron [µg/dl] | 174.1 ± 5.58 | 184.5 ± 12.04 | 165.1 ± 8.49 | 140.4 ± 12.42 | n.s. | p<0.05 | p<0.05 | n.s. | n.s. |

| Table 21: Clinical-chemical parameters, 2nd sample. Data are presented as mean ± standard error of mean. | | | | | | | | | |
|---|---|---|---|---|---|---|---|---|---|
| Parameter | Control (A) | | Mutant (B) | | A~B | A~B | ANOVA | | |
| | Male | Female | Male | Female | Male | Female | genotype | sex | inter. |
| | (n=11) | (n=9) | (n=9) | (n=11) | *p-value* | *p-value* | *p-value* | *p-value* | *p-value* |
| Potassium [mmol/l] | 3.98 ± 0.09 | 3.71 ± 0.09 | 4.03 ± 0.07 | 3.98 ± 0.07 | n.s. | p<0.05 | p<0.05 | p<0.05 | n.s. |
| Total Protein [g/dl] | 5.56 ± 0.06 | 5.02 ± 0.05 | 5.33 ± 0.07 | 4.98 ± 0.05 | p<0.05 | n.s. | p<0.05 | p<0.001 | n.s. |
| Albumin [g/dl] | 2.8 ± 0.038 | 2.622 ± 0.04 | 2.6 ± 0.047 | 2.455 ± 0.039 | p<0.01 | p<0.01 | p<0.001 | p<0.001 | n.s. |
| Triglycerides [mg/dl] | 306 ± 20.4 | 251 ± 20.1 | 288 ± 18.1 | 220 ± 13.5 | n.s. | n.s. | n.s. | p<0.01 | n.s. |
| ASAT [U/l] | 45.5 ± 9.57 | 40.9 ± 4.2 | 41.6 ± 2.53 | 80.7 ± 24.55 | n.s. | n.s. | n.s. | n.s. | n.s. |
| TIBC [µg/dl] | 469.8 ± 10.34 | 481.5 ± 7.57 | 434.1 ± 15.91 | 450.2 ± 7.58 | n.s. | p<0.01 | p<0.01 | n.s. | n.s. |
| Ferritin [ng/ml] | 18.2 ± 1.15 | 21.5 ± 1.02 | 16.5 ± 1.41 | 19.5 ± 0.96 | n.s. | n.s. | n.s. | p<0.01 | n.s. |
| Transferrin [mg/dl] | 137.1 ± 2.14 | 131.8 ± 1.52 | 130.6 ± 2.07 | 128.3 ± 0.99 | p<0.05 | n.s. | p<0.01 | p<0.05 | n.s. |
| Iron [µg/dl] | 174.4 ± 2.92 | 178.3 ± 7.84 | 147.4 ± 3.38 | 169.6 ± 4.03 | p<0.001 | n.s. | p<0.001 | p<0.01 | n.s. |

Table 22: Hematological parameters, 1st sample.
Data are presented as mean ± standard error of mean.

| Parameter | Control (A) | | Mutant (B) | | A~B | A~B | ANOVA | | |
|---|---|---|---|---|---|---|---|---|---|
| | Male | Female | Male | Female | Male | Female | geno-type | sex | inter. |
| | (n=11) | (n=9) | (n=9) | (n=11) | *p-value* | *p-value* | *p-value* | *p-value* | *p-value* |
| WBC [10³/µl] | 6.7 ± 0.32 | 9 ± 0.85 | 10.1 ± 0.51 | 10.5 ± 0.65 | p<0.001 | n.s. | p<0.01 | p<0.05 | p<0.05 |
| RBC [10⁶/µl] | 9.7 ± 0.08 | 9.3 ± 0.08 | 12.5 ± 0.17 | 12 ± 0.07 | p<0.001 | p<0.001 | p<0.001 | n.s. | n.s. |
| PLT [10³/µl] | 998 ± 30.8 | 1059 ± 34.4 | 885 ± 32 | 863 ± 28.4 | p<0.05 | p<0.001 | p<0.001 | n.s. | n.s. |
| Hemoglobin [g/dl] | 14.2 ± 0.48 | 15 ± 0.13 | 16.8 ± 0.26 | 16.7 ± 0.12 | p<0.001 | p<0.001 | p<0.001 | n.s. | n.s. |
| Hematocrit [%] | 53.2 ± 0.46 | 52.1 ± 0.4 | 60.8 ± 0.82 | 59.2 ± 0.35 | p<0.001 | p<0.001 | p<0.001 | n.s. | n.s. |
| MCV [fl] | 54.9 ± 0.21 | 56 ± 0.17 | 48.7 ± 0.17 | 49.3 ± 0.14 | p<0.001 | p<0.001 | p<0.001 | n.s. | n.s. |
| MCH [pg] | 14.7 ± 0.49 | 16.2 ± 0.06 | 13.5 ± 0.11 | 13.9 ± 0.03 | p<0.05 | p<0.001 | p<0.001 | p<0.05 | n.s. |
| MCHC [g/dl] | 26.8 ± 0.89 | 28.9 ± 0.11 | 27.7 ± 0.22 | 28.2 ± 0.05 | n.s. | p<0.001 | n.s. | p<0.05 | n.s. |
| RDW [% of MCV] | 12.4 ± 0.06 | 12.1 ± 0.06 | 12.3 ± 0.08 | 11.8 ± 0.09 | n.s. | p<0.05 | p<0.01 | p<0.001 | n.s. |
| MPV [fl] | 6.15 ± 0.12 | 5.97 ± 0.03 | 6.09 ± 0.08 | 5.93 ± 0.09 | n.s. | n.s. | n.s. | n.s. | n.s. |

Table 23: Hematological parameters, 2nd sample.
Data are presented as mean ± standard error of mean.

| Parameter | Control (A) | | Mutant (B) | | A~B | A~B | ANOVA | | |
|---|---|---|---|---|---|---|---|---|---|
| | Male | Female | Male | Female | Male | Female | geno-type | sex | inter. |
| | (n=10) | (n=10) | (n=10) | (n=10) | *p-value* | *p-value* | *p-value* | *p-value* | *p-value* |
| WBC [10³/µl] | 7 ± 0.47 | 8.1 ± 0.32 | 8.2 ± 0.71 | 9.8 ± 0.69 | n.s. | p<0.05 | p<0.05 | p<0.05 | n.s. |
| RBC [10⁶/µl] | 9.3 ± 0.09 | 9.2 ± 0.13 | 12.1 ± 0.12 | 11.9 ± 0.14 | p<0.001 | p<0.001 | p<0.001 | n.s. | n.s. |
| PLT [10³/µl] | 1436 ± 37.7 | 1270 ± 55 | 1105 ± 43.5 | 1007 ± 40.2 | p<0.001 | p<0.01 | p<0.001 | p<0.01 | n.s. |
| Hemoglobin [g/dl] | 15.1 ± 0.16 | 15 ± 0.22 | 16.6 ± 0.13 | 16.4 ± 0.13 | p<0.001 | p<0.001 | p<0.001 | n.s. | n.s. |
| Hematocrit [%] | 52.2 ± 0.55 | 52.2 ± 0.93 | 58.7 ± 0.49 | 58.1 ± 0.62 | p<0.001 | p<0.001 | p<0.001 | n.s. | n.s. |
| MCV [fl] | 56.2 ± 0.23 | 56.3 ± 0.33 | 48.4 ± 0.18 | 48.9 ± 0.16 | p<0.001 | p<0.001 | p<0.001 | n.s. | n.s. |
| MCH [pg] | 16.3 ± 0.09 | 16.2 ± 0.11 | 13.7 ± 0.07 | 13.8 ± 0.09 | p<0.001 | p<0.001 | p<0.001 | n.s. | n.s. |
| MCHC [g/dl] | 29 ± 0.1 | 28.7 ± 0.14 | 28.3 ± 0.09 | 28.3 ± 0.11 | p<0.001 | p<0.05 | p<0.001 | n.s. | n.s. |
| RDW [% of MCV] | 13.5 ± 0.07 | 14 ± 0.22 | 12.8 ± 0.08 | 13.4 ± 0.08 | p<0.001 | p<0.05 | p<0.001 | p<0.001 | n.s. |
| MPV [fl] | 5.8 ± 0.07 | 6 ± 0.12 | 6.01 ± 0.12 | 6.04 ± 0.08 | n.s. | n.s. | n.s. | n.s. | n.s. |

#### Immunology

Results FACS analysis and plasma immunoglobulin levels determined from blood sample collected at age 20 weeks from *ad libitum* fed mice

**Table 24: Basic parameters analyzed in the Immunology Screen by Flow Cytometry**
Frequencies of main leukocyte subsets in blood of male and female MVD013 transgenic and wild-type mice from second bleeding [% of CD45+viable leukocytes, respectively of parent gate].

| Variable | Control (A) | | | Mutant (B) | | | A ~ B | |
|---|---|---|---|---|---|---|---|---|
| | Male | Female | | Male | Female | | Male | Female |
| | 10 | 10 | *p - value* | 10 | 10 | *p - value* | *p - value* | *p - value* |
| 45+/CD3+ | 27.8 ± 0.97 | 29.5 ± 1.25 | n.s. | 25.8 ± 1.36 | 22.5 ± 1.51 | n.s. | n.s. | p<0.01 |
| 45+/CD3+ CD4+ | 16.67 ± 0.78 | 16.34 ± 0.8 | n.s. | 14.47 ± 1.07 | 11.54 ± 0.84 | p<0.05 | n.s. | p<0.001 |
| 45+/CD3+ CD8+ | 10.45 ± 0.26 | 12.03 ± 0.48 | p<0.05 | 10.16 ± 0.58 | 9.74 ± 0.63 | n.s. | n.s. | p<0.01 |
| CD3+4+/ CD25+ | 3.1 ± 0.25 | 3.8 ± 0.14 | p<0.05 | 3.4 ± 0.19 | 4.9 ± 0.33 | p<0.01 | n.s. | p<0.01 |
| 45+/gamma delta TCR+CD3+ | 1.71 ± 0.21 | 1.39 ± 0.16 | n.s. | 1.09 ± 0.14 | 1.04 ± 0.1 | n.s. | p<0.05 | n.s. |
| 4+3+/62L+ | 24.97 ± 3.07 | 18.73 ± 1.75 | n.s. | 20.27 ± 2.82 | 17.94 ± 1.14 | n.s. | n.s. | n.s. |
| 4+3+/44L+ | 57.9 ± 0.676 | 56.42 ± 1.03 | n.s. | 59.62 ± 1.335 | 58.17 ± 0.92 | n.s. | n.s. | n.s. |
| 8+3+/62L+ | 47 ± 4.01 | 39.4 ± 2.56 | n.s. | 40.4 ± 3.59 | 37.2 ± 1.75 | n.s. | n.s. | n.s. |
| 8+3+/44+ | 45.99 ± 1.08 | 44.61 ± 1.35 | n.s. | 48.43 ± 1.22 | 47.5 ± 1.3 | n.s. | n.s. | n.s. |
| 45+/Gr1+ CD11b+ | 25 ± 2.43 | 19.4 ± 1.14 | n.s. | 32.3 ± 3.58 | 35 ± 2.01 | n.s. | n.s. | p<0.001 |
| 45+/nonGrno nNKCD11b+ | 7 ± 0.4 | 7 ± 0.3 | n.s. | 7 ± 0.5 | 9 ± 0.8 | n.s. | n.s. | p<0.05 |
| 45+/CD19+ | 28.1 ± 1.27 | 37 ± 1.57 | p<0.001 | 25.1 ± 1.84 | 25.9 ± 0.6 | n.s. | n.s. | p<0.001 |
| CD19+/IgD+ | 89.8 ± 0.55 | 91.8 ± 0.66 | p<0.05 | 88.9 ± 0.65 | 87.9 ± 0.42 | n.s. | n.s. | p<0.001 |
| CD19+/ CD5+ | 3.21 ± 0.2 | 3.71 ± 0.2 | n.s. | 3.24 ± 0.3 | 3.02 ± 0.2 | n.s. | n.s. | p<0.05 |
| CD19+/B220+ MHC-II+ | 42.9 ± 1.5 | 50.8 ± 1.6 | p<0.01 | 44.9 ± 2 | 49.7 ± 1.8 | n.s. | n.s. | n.s. |

**Table 26: Basic parameters analyzed in the Immunology Screen by Bioplex/ELISA**
Concentration [µg/ml] of antibodies of different isotypes, and levels of autoantibodies (OD) in blood plasma from mutant and control mice

| Parameter | Control (A) | | | Mutant (B) | | | A ~ B | |
|---|---|---|---|---|---|---|---|---|
| | Male | Female | | Male | Female | | Male | Female |
| | 10 | 10 | *p - value* | 10 | 10 | *p - value* | *p - value* | *p - value* |
| IgM | 560.8 ± 34.38 | 830.2 ± 69.3 | p<0.01 | 650.5 ± 24.3 | 869.4 ± 139.37 | n.s. | p<0.05 | n.s. |
| IgA | 676.29 ± 85.87 | 947.45 ± 101.32 | n.s. | 1220.2 ± 220.48 | 1355.72 ± 123.51 | n.s. | p<0.05 | p<0.05 |
| IgG3 | 483.85 ± 79 | 552.1 ± 78.12 | n.s. | 594.17 ± 54.97 | 1047.34 ± 167.76 | p<0.05 | n.s. | p<0.05 |
| IgG1 | 296.5 ± 25.95 | 543.9 ± 65.91 | p<0.01 | 390.3 ± 25.25 | 494.9 ± 82.22 | n.s. | p<0.05 | n.s. |
| IgG2a | 756.62 ± 113.81 | 1259.42 ± 186.5 | p<0.05 | 1523.62 ± 157.05 | 1931.81 ± 129.27 | n.s. | p<0.01 | p<0.05 |
| IgG2b | 121.21 ± 8.65 | 465.69 ± 79.81 | p<0.01 | 373.77 ± 41.93 | 718.81 ± 54.14 | p<0.001 | p<0.001 | p<0.05 |
| Anti- D N A | 0.67 ± 0.104 | 0.636 ± 0.08 | n.s. | 0.738 ± 0.061 | 0.732 ± 0.054 | n.s. | n.s. | n.s. |
| Rheumatoid factor | 0.1 ± 0.02 | 0.1 ± 0.01 | n.s. | 0.1 ± 0.01 | 0.1 ± 0.02 | n.s. | n.s. | n.s. |

#### Steroid screen (age 23 weeks)

**Sample preparation.** Since no steroid ELISA kits are available for mouse samples, human ELISA kits have been adopted. Prior to measurement, the steroids have to be extracted from the matrix by liquid/liquid-extraction to avoid mouse plasma matrix effects. 75 µl of plasma were extracted three times in each case with a tenfold excess of *tert*-butylmethylether (TBME). The combined organic extracts were evaporated, dissolved *de novo* in TBME, subdivided equally for the two ELISA tests (DHEA and testosterone) and evaporated again. The material was reconstituted for the respective kit, DHEA in assay buffer, testosterone in steroid free serum.

**ELISA.** The steroids were quantified by competitive ELISA according to the manufacturer's protocols. The plates were read in a standard microplate reader at a wavelength of 405 nm (DHEA) and 450 nm (testosterone). The concentrations were calculated upon the calibration from the respective standard curve and reported in pg/ml (DHEA) and ng/ml (testosterone). The sensitivity of the tests is 2.9 pg/ml for DHEA and 0.083 ng/ml for testosterone.

We used the following ELISA kits:

Testosterone ELISA: DRG Instruments GmbH, Catalog No. EIA-1559

DHEA ELISA: AssayDesigns, Catalog No. 901-093

Table 28: Plasma levels of DHEA and testosterone of *MVD013* mice (22 weeks old)
Data are presented as median (25 %/75 % - interquartile range)

| | Control (A) | | | Mutant (B) | | | A~B | A~B |
|---|---|---|---|---|---|---|---|---|
| | Male | Female | | Male | Female | | Male | Female |
| | (n=11) | (n=9) | *p - value* | (n=9) | (n=11) | *p - value* | *p - value* | *p - value* |
| DHEA [pg/ml] | 19.2 (7.1/ 37.0) | 66.5 (43.9 / 83.0) | 0.002 | 23.6 (<2.9 / 50.0) | 20.4↓ (3.9 / 56.3) | n.s. | n.s. | 0.012 |
| Testosterone [ng/ml] | 0.624 (0.365 / 4.659) | 0.108 (0.089 / 0.131) | 0.002 | 0.733 (0.423 / 1.338) | 0.102 (<0.083 / 0.175) | 0.002 | n.s. | n.s. |

#### Cardio—vascular Screen

**Tail-cuff blood pressure measurement:** Blood pressure was measured in unanesthetized mice with a non-invasive tail-cuff method using the MC4000 Blood Pressure Analysis Systems (Hatteras Instruments Inc., Cary, North Carolina, USA). Four animals were restrained on a pre-warmed metal platform in metal boxes. The tails were looped through a tail-cuff and fixed in a notch containing an optical path with a LED light and a photosensor.

The blood pulse wave in the tail artery is detected as transformed into an optical pulse signal by measurement of light extinction. Pulse detection, cuff inflation and pressure evaluation are automated by the system software. After five initial inflation runs for habituation, 12 measurement runs are performed for each animal in one session. Runs with movement artifacts are excluded.

After one day of training, in which the animals are habituated to the apparatus and protocol, the measurements are performed on four consecutive days between 8:30 and 11:30 AM.

| Table 29: Blood Pressure Parameters<br>Data are presented as mean +/- standard error of mean. | | | | | | | | | |
|---|---|---|---|---|---|---|---|---|---|
| Parameter | Control (A) | | Mutant (B) | | ANOVA | | | Post hoc test | |
| | | | | | | | | A~B | A~B |
| | Male | Female | Male | Female | Sex | Genotype | Interact. | Male | Female |
| | ( n = 10 ) | ( n = 10 ) | ( n = 10 ) | ( n = 10 ) | *p - value* | *p - value* | *p - value* | *p - value* | *p - value* |
| Systolic pressure [mm Hg] | 112.1 +/- 1.6 | 117.6 +/- 3.5 | 109.7 +/- 3.9 | 111.1 +/- 5.2 | n.s. | n.s. | n.s. | n.s. | n.s. |
| Diastolic pressure [mm Hg] | 101.9 +/- 1.8 | 109.0 +/- 4.0 | 97.0 +/- 3.8 | 97.9 +/- 4.9 | n.s. | p<0.05 | n.s. | n.s. | p=0.058 |
| Mean arterial pressure [mm Hg] | 104.9 +/- 1.7 | 111.5 +/- 3.8 | 100.9 +/- 3.8 | 102.0 +/- 4.9 | n.s. | n.s. | n.s. | n.s. | n.s. |
| Pulse [bpm] | 527.7 +/- 17.1 | 555.0 +/- 6.4 | 507.1 +/- 10.6 | 529.9 +/- 15.4 | n.s. | n.s. | n.s. | n.s. | n.s. |

**Echocardiography:** Left ventricular function was determined using a small animal ultrasound biomicroscope with a 30-MHz transducer and 30 Hz frame rate (Vevo 660; VisualSonics, Toronto, Ontario). The shaved and anesthetized mice (1% isoflurane inhalation, Baxter, Munich, Germany) were fixed in supine position on a heated platform equipped with ECG electrodes for heart rate monitoring. Body temperature was maintained at 36–38°C, monitored via a rectal thermometer (Indus Instruments, Houston, Texas, USA). Left ventricular parasternal short-axis views were obtained at the papillary muscle level recording 2-dimensional B-Mode images and time-motion M-mode images.

| Table 30: Echocardiography<br>Data are presented as mean +/- standard error of mean. | | | | | | | | | |
|---|---|---|---|---|---|---|---|---|---|
| Parameter | Control (A) | | Mutant (B) | | ANOVA | | | Post hoc test | |
| | | | | | | | | A~B | A~B |
| | Male | Female | Male | Female | Sex | Genotype | Interact. | Male | Female |
| | ( n = 10 ) | ( n = 7 ) | ( n = 7 ) | ( n = 8 ) | *p - value* | *p - value* | *p - value* | *p - value* | *p - value* |
| LVID dia [mm] | 4.18 +/- 0.07 | 3.81 +/- 0.07 | 3.99 +/- 0.10 | 3.77 +/- 0.07 | p<0.001 | n.s. | n.s. | n.s. | n.s. |
| LVID sys [mm] | 3.09 +/- 0.11 | 2.65 +/- 0.13 | 2.85 +/- 0.14 | 2.61 +/- 0.08 | p<0.01 | n.s. | n.s. | n.s. | n.s. |
| Heart rate [bpm] | 483 +/- 13 | 453 +/- 14 | 436 +/- 15 | 465 +/- 15 | n.s. | p=0.055 | n.s. | p<0.05 | n.s. |
| Fractional shortening [%] | 26.2 +/- 1.8 | 30.6 +/- 2.2 | 28.7 +/- 2.0 | 30.9 +/- 0.9 | n.s. | n.s. | n.s. | n.s. | n.s. |
| LV volume dia [ml] | 77.9 +/- 2.9 | 62.4 +/- 2.9 | 70.0 +/- 4.0 | 61.0 +/- 2.7 | p<0.001 | n.s. | n.s. | n.s. | n.s. |
| LV volume sys [ml] | 38.5 +/- 3.4 | 26.5 +/- 3.1 | 31.7 +/- 3.5 | 25.1 +/- 1.8 | p<0.01 | n.s. | n.s. | n.s. | n.s. |
| Ejection Fraction [%] | 51.3 +/- 2.8 | 58.4 +/- 3.2 | 55.4 +/- 3.0 | 59.2 +/- 1.4 | n.s. | n.s. | n.s. | n.s. | n.s. |

LV – left ventricle, LVID – left ventricle inner diameter

**Heart Weight Determination:** During the final examination in the Pathology Screen the heart weight was determined together with body weight and tibia length. Briefly, mice were sacrificed by $CO_2$ inhalation, weighed and opened from the ventral midline. Exsanguination was achieved by cutting the dorsal aorta. Prior to dissection the heart was inspected for abnormalities or excessive fat. For excision the heart was removed from the pericardial membrane and the major vessels were cut through at the point they enter or exit the atria. The heart weight was obtained wet after blotting the organ on paper towels. The tibia length was determined from the left tibia of the mouse using a ruler.

| Table 31: Heart weight<br>Data are presented as mean +/- standard error of mean. | | | | | | | | | |
|---|---|---|---|---|---|---|---|---|---|
| Parameter | Control (A) | | Mutant (B) | | ANOVA | | | Post hoc test | |
| | | | | | | | | A~B | A~B |
| | Male | Female | Male | Female | Sex | Genotype | Interact. | Male | Female |
| | ( n = 11 ) | ( n = 9 ) | ( n = 9 ) | ( n = 11 ) | *p - value* | *p - value* | *p - value* | *p - value* | *p - value* |
| Body Weight [g] | 32.5 +/- 0.5 | 27.9 +/- 1.3 | 29.4 +/- 0.6 | 26.9 +/- 0.5 | p<0.001 | p<0.01 | n.s. | p<0.01 | n.s. |
| Heart Weight [g] | 0.14 +/- 0.00 | 0.12 +/- 0.00 | 0.13 +/- 0.00 | 0.12 +/- 0.00 | p<0.001 | p<0.001 | p<0.01 | p<0.001 | n.s. |
| Tibia lenght [mm] | 16.5 +/- 0.2 | 16.3 +/- 0.2 | 16.6 +/- 0.2 | 16.4 +/- 0.2 | n.s. | n.s. | n.s. | n.s. | n.s. |
| HW/TBL | 8.81 +/- 0.21 | 7.16 +/- 0.13 | 7.77 +/- 0.12 | 7.04 +/- 0.10 | p<0.001 | p<0.001 | p<0.01 | p<0.001 | n.s. |
| HW/BW | 4.46 +/- 0.10 | 4.25 +/- 0.18 | 4.39 +/- 0.11 | 4.29 +/- 0.08 | n.s. | n.s. | n.s. | n.s. | n.s. |

HW – heart weight, TBL – tibial length, BW – body weight

#### Lung function Screen (age 23 weeks)

**Whole Body Plethysmography:** A commercially available system from Buxco® Electronics (Sharon, Connecticut) was used to assess breathing patterns in unrestrained animals according to the principle described by Drorbaugh and Fenn (1955). It measures the pressure changes which arise from inspiratory and expiratory temperature and humidity fluctuations during breathing

| 2. Respiratory rate and timing at sleep, rest and activity | Data are presented as mean ± standard error of mean. | | | | | | | |
|---|---|---|---|---|---|---|---|---|
| **Parameter** | **Control (A)** | | | **Mutant (B)** | | | **A~B** | **A~B** |
| | **Male** | **Female** | | **Male** | **Female** | | **Male** | **Female** |
| | **(n=6)** | **(n=6)** | ***p - value*** | **(n=6)** | **(n=5)** | ***p - value*** | ***p - value*** | ***p - value*** |
| **Sleep** | | | | | | | | |
| **f [1/min]** | 147.6 ± 4.8 | 144.7 ± 4.6 | n.s. | 145.1 ± 2.0 | 123.3 ± 1.0 | < 0.001 | n.s. | < 0.01 |
| **Ti [ms]** | 124.5 ± 3.2 | 137.0 ± 3.9 | < 0.05 | 133.4 ± 3.6 | 147.4 ± 4.1 | < 0.05 | n.s. | n.s. |
| **Te [ms]** | 284.2 ± 11.2 | 279.8 ± 10.5 | n.s. | 280.3 ± 3.9 | 339.3 ± 5.5 | < 0.001 | n.s. | < 0.01 |
| **Ti/TT** | 0.31 ± 0.01 | 0.33 ± 0.01 | < 0.05 | 0.32 ± 0.01 | 0.30 ± 0.01 | n.s. | n.s. | < 0.05 |
| **Rest** | | | | | | | | |
| **f [1/min]** | 295.7 ± 4.3 | 283.4 ± 3.8 | n.s. | 295.8 ± 4.1 | 287.8 ± 2.8 | n.s. | n.s. | n.s. |
| **Ti [ms]** | 63.7 ± 1.6 | 68.4 ± 1.1 | < 0.05 | 64.6 ± 1.3 | 63.5 ± 1.1 | n.s. | n.s. | < 0.02 |
| **Te [ms]** | 139.4 ± 2.0 | 143.5 ± 3.3 | n.s. | 138.4 ± 2.4 | 145.0 ± 2.3 | n.s. | n.s. | n.s. |
| **Ti/TT** | 0.31 ± 0.01 | 0.32 ± 0.01 | n.s. | 0.32 ± 0.01 | 0.30 ± 0.01 | n.s. | n.s. | n.s. |
| **Activity** | | | | | | | | |
| **f [1/min]** | 487.5 ± 2.7 | 462.8 ± 7.3 | < 0.01 | 480.7 ± 5.9 | 456.9 ± 7.0 | < 0.05 | n.s. | n.s. |
| **Ti [ms]** | 40.4 ± 0.3 | 42.3 ± 0.5 | < 0.02 | 41.5 ± 0.4 | 41.0 ± 0.6 | n.s. | n.s. | n.s. |
| **Te [ms]** | 82.7 ± 0.5 | 87.5 ± 1.6 | < 0.02 | 83.4 ± 1.3 | 90.4 ± 1.6 | < 0.01 | n.s. | n.s. |
| **Ti/TT** | 0.33 ± 0.00 | 0.33 ± 0.00 | n.s. | 0.33 ± 0.00 | 0.31 ± 0.00 | < 0.01 | n.s. | < 0.01 |

| 3. Tidal volume and flow rates at sleep, rest and activity Data are presented as mean ± standard error of mean. | | | | | | | | |
|---|---|---|---|---|---|---|---|---|
| Parameter | Control (A) | | | Mutant (B) | | | A~B | A~B |
| | Male | Female | | Male | Female | | Male | Female |
| | (n=6) | (n=6) | *p - value* | (n=6) | (n=5) | *p - value* | *p - value* | *p - value* |
| Sleep | | | | | | | | |
| TV [ml] | 0.25 ± 0.01 | 0.28 ± 0.01 | n.s. | 0.27 ± 0.01 | 0.28 ± 0.01 | n.s. | n.s. | n.s. |
| PIF [ml/s] | 3.3 ± 0.1 | 3.3 ± 0.1 | n.s. | 3.2 ± 0.1 | 3.1 ± 0.1 | n.s. | n.s. | n.s. |
| PEF [ml/s] | 2.1 ± 0.2 | 2.3 ± 0.2 | n.s. | 2.3 ± 0.1 | 2.1 ± 0.1 | n.s. | n.s. | n.s. |
| MIF [ml/s] | 2.0 ± 0.0 | 2.0 ± 0.0 | n.s. | 2.0 ± 0.0 | 1.9 ± 0.0 | n.s. | n.s. | n.s. |
| MEF [ml/s] | 0.88 ± 0.02 | 0.99 ± 0.04 | n.s. | 0.95 ± 0.02 | 0.84 ± 0.03 | n.s. | n.s. | < 0.02 |
| Rest | | | | | | | | |
| TV [ml] | 0.20 ± 0.01 | 0.21 ± 0.01 | n.s. | 0.20 ± 0.00 | 0.19 ± 0.01 | n.s. | n.s. | n.s. |
| PIF [ml/s] | 5.6 ± 0.1 | 5.4 ± 0.2 | n.s. | 5.5 ± 0.2 | 5.5 ± 0.2 | n.s. | n.s. | n.s. |
| PEF [ml/s] | 3.3 ± 0.3 | 3.4 ± 0.2 | n.s. | 3.2 ± 0.1 | 3.2 ± 0.2 | n.s. | n.s. | n.s. |
| MIF [ml/s] | 3.1 ± 0.1 | 3.0 ± 0.1 | n.s. | 3.1 ± 0.1 | 3.0 ± 0.1 | n.s. | n.s. | n.s. |
| MEF [ml/s] | 1.4 ± 0.0 | 1.4 ± 0.1 | n.s. | 1.5 ± 0.0 | 1.3 ± 0.1 | n.s. | n.s. | n.s. |
| Activity | | | | | | | | |
| TV [ml] | 0.20 ± 0.01 | 0.19 ± 0.01 | n.s. | 0.20 ± 0.01 | 0.18 ± 0.00 | < 0.01 | n.s. | n.s. |
| PIF [ml/s] | 8.4 ± 0.3 | 8.1 ± 0.4 | n.s. | 8.4 ± 0.3 | 7.5 ± 0.2 | n.s. | n.s. | n.s. |
| PEF [ml/s] | 5.1 ± 0.3 | 5.0 ± 0.4 | n.s. | 5.1 ± 0.2 | 4.5 ± 0.2 | n.s. | n.s. | n.s. |
| MIF [ml/s] | 4.9 ± 0.2 | 4.6 ± 0.2 | n.s. | 4.9 ± 0.1 | 4.3 ± 0.1 | < 0.02 | n.s. | n.s. |
| MEF [ml/s] | 2.4 ± 0.1 | 2.2 ± 0.1 | n.s. | 2.5 ± 0.1 | 2.0 ± 0.1 | < 0.01 | n.s. | n.s. |

| 4. Minute ventilation and body size/weight related parameters at sleep, rest and activity Data are presented as mean ± standard error of mean. | | | | | | | | |
|---|---|---|---|---|---|---|---|---|
| Parameter | Control (A) | | | Mutant (B) | | | A~B | A~B |
| | Male | Female | | Male | Female | | Male | Female |
| | (n=6) | (n=6) | *p - value* | (n=6) | (n=5) | *p - value* | *p - value* | *p - value* |
| Sleep | | | | | | | | |
| sTV [µl/g] | 8.0 ± 0.4 | 10.0 ± 0.7 | < 0.02 | 8.2 ± 0.3 | 11.4 ± 0.1 | < 0.001 | n.s. | n.s. |
| MV [ml/min] | 36.1 ± 0.8 | 39.1 ± 1.5 | n.s. | 37.9 ± 0.8 | 34.1 ± 0.8 | n.s. | n.s. | < 0.05 |
| sMV [ml/min/g] | 1.1 ± 0.1 | 1.4 ± 0.1 | < 0.05 | 1.2 ± 0.0 | 1.4 ± 0.0 | < 0.001 | n.s. | n.s. |
| Rest | | | | | | | | |
| sTV [µl/g] | 6.3 ± 0.2 | 7.5 ± 0.6 | n.s. | 6.2 ± 0.1 | 7.8 ± 0.3 | < 0.001 | n.s. | n.s. |
| MV [ml/min] | 54.8 ± 0.8 | 53.4 ± 2.7 | n.s. | 55.7 ± 1.8 | 51.0 ± 2.0 | n.s. | n.s. | n.s. |
| sMV [ml/min/g] | 1.7 ± 0.1 | 1.9 ± 0.2 | n.s. | 1.7 ± 0.0 | 2.1 ± 0.1 | n.s. | n.s. | n.s. |
| Activity | | | | | | | | |
| sTV [µl/g] | 6.3 ± 0.2 | 7.0 ± 0.5 | n.s. | 6.3 ± 0.2 | 7.1 ± 0.3 | n.s. | n.s. | n.s. |
| MV [ml/min] | 95.3 ± 3.8 | 87.2 ± 3.9 | n.s. | 96.4 ± 3.1 | 78.8 ± 2.9 | < 0.01 | n.s. | n.s. |
| sMV [ml/min/g] | 3.0 ± 0.1 | 3.2 ± 0.2 | n.s. | 3.0 ± 0.1 | 3.2 ± 0.2 | n.s. | n.s. | n.s. |

Supplementary Table 1: Overview of macroscopic and histologic pathology results

| Phenotype Age<br>Number of animals | ICC hyperplasia stomach | ICC hyperplasia cecum/colon | Enlarged GALT cecum/colon | Mamma tumor<br>**Metastasis** | Liver tumor | Other findings | Description |
|---|---|---|---|---|---|---|---|
| Mut<br><2 months<br>N=3 | 1/3 f<br>(+) | 1/3 f<br>(+) | 1/3 f<br>(+) | 0/3 f | 0/3 f | 0/3 f | |
| Mut<br>5-12 months<br>N=24 | 14/14 f<br>(+ or ++)<br>10/10 m<br>(+ or ++) | 14/14 f<br>(+ or ++)<br>8/10 m<br>(+ or ++) | 13/14 f<br>(+ or ++)<br>7/10 m<br>(+ or ++) | 4/14 f<br>0/10m | 0/14 f<br>0/10 m | 6/14 f<br>5/10 m | Constipation with megacecum 10/24, lymphoma 1/24 |
| Mut<br>12-15 months<br>N=35 | 17/17 f<br>(+ or ++)<br>18/18 m<br>(+ or ++) | 16/17 f<br>(+ or ++)<br>18/18 m<br>(+ or ++) | 14/17 f<br>(+/++/ +++)<br>intestine of remaining animals too dilated to see GALT<br>18/18 m<br>(+ or ++) | 14/17 f<br>**6/17 f**<br>0/18 m | 0/12 f<br>1/18 m | 17/17 f<br>3/17 f<br>18/18 m | Constipation with megacecum in all animals<br>leukosis 1f,<br>liver necrosis 2f |
| Mut<br>15 – 19 months<br>N=14 | 14/14 m<br>(+/++/+++) | 13/14 m<br>(+) | 12/14 m<br>(++ or +++) | 0/14 m | 1/14 m | 12/14 m | Constipation with megacecum |
| WT<br>5-12 months<br>N=14 | 0/6 f<br>0/8 m | 0/6 f<br>0/8 m | 3/6 f<br>(+ or ++)<br>5/8 m<br>(+ or ++) | 0/6 f<br>0/8 m | 0/6 f<br>0/8 m | 1/6 f<br>0/8 m | Cystic mass ovary |
| WT<br>12 -20 months<br>N=33 | 0/13 f<br>0/20 m | 0/13 f<br>0/20 m | 5/13 f<br>6/20 m<br>(+/ ++/+++) | 0/13 f<br>0/20 m | 0/13 f<br>6/20 m | 2/13 f<br>0/13 m | Cystic mass ovary |

F – female, m – male, +/++/+++ - semiquantitative grading of severity

#### Supplement 2:
#### Analysis of bone marrow of KitN824K/WT and control mice

#### Methods

**Apoptosis assay:** Bone marrow was isolated from femurs by flushing. $5x10^5$ cells were transferred into a 1,5mL Eppendorf tube and centrifuged (13.000 rpm, 15sec). Cell pellets were washed with 1,5 mL Annexin-V-binding buffer. The supernatant was aspirated and the cells were resuspended in 100µL Annexin-V-binding buffer. Then 4µl of Annexin-V-AF647 (BioLegend, 640911) and 1,5µl (3µM) SytoxOrange staining solution (Molecular Probes S11368) were pipetted into the Eppendorf tube. To detect apoptosis in the Kit+/Sca-1+/Lin- (KSL) and Kit+/Sca-1-/Lin- (KL) bone marrow population 1µL of each antibody were added (anti lineage cocktail FITC (Biolegend, 78022), anti sca-1 Pacific Blue (Biolegend, 108120), anti c-kit APC-eFluor780 (eBioScience 471171-82), gently vortexed and incubated 20-30 min. at room temperature in the dark. The cells were washed once with Annexin-V-binding buffer. Pellets were resuspended in 250 µl Annexin-V-binding buffer and analyzed on a FACS Canto Cytometer (Becton Dickinson).

**Colony assays:** BM cells were seeded in methylcellulose media with (M3334) or without Epo (M3234) (STEMCELL Technologies) for CFU-E assays or in media containing SCF, IL-3, and IL6, with EPO (M3434) (STEMCELL Technologies) for BFU-E, CFU-GM, and CFU-GEMM colony growth and incubated for 4–8 d at 37°C, 5% CO2. CFU-E colony growth was scored on day 4, and BFU-E, CFU-GM, and CFU-GEMM colony growth was scored on day 8 after benzidine staining.

#### Results

Bone marrow derived from the femurs of six months old *Kit*$^{N824K/WT}$ mutant mice and age matched wild-type littermates contained similar numbers of Kit-expressing stem cells including comparable percentages of common myeloid progenitor cells (CMPs) and megacaryocytic-erythroid progenitor cells (MEPs) in both genotypes. Also apoptosis rates in the different stem cell populations were not significantly different at this age (Fig. S1 A,B,C). In bone marrow collected from *Kit*$^{N824/WT}$ animals at one year the proportion of Kit+ stem cells was increased and included a reduced portion of CMPs and an increased percentage of MEPs compared to bone. Surprisingly, the proportion of apoptotic stem cells was clearly increased at this age in mutants, for all stem cell populations investigated (Fig. S1 D,E,F). Colony assay analysis, however, revealed an increased proliferative activity especially of cells from the erythroid (E) and granulocytic-macrophage (GM) lineage, with increased CFU-GM counts and CFU-E counts independent of erythropoietin stimulation and similar BFU-E numbers in mutants compared to controls at the age of six months (Fig. S1 G,H). Bone marrow derived from one year old mutant animals in contrast displayed decreased CFU-E, BFU-E and CFU-GM counts compared to controls, suggesting decreased proliferative activity (S1 I,J).

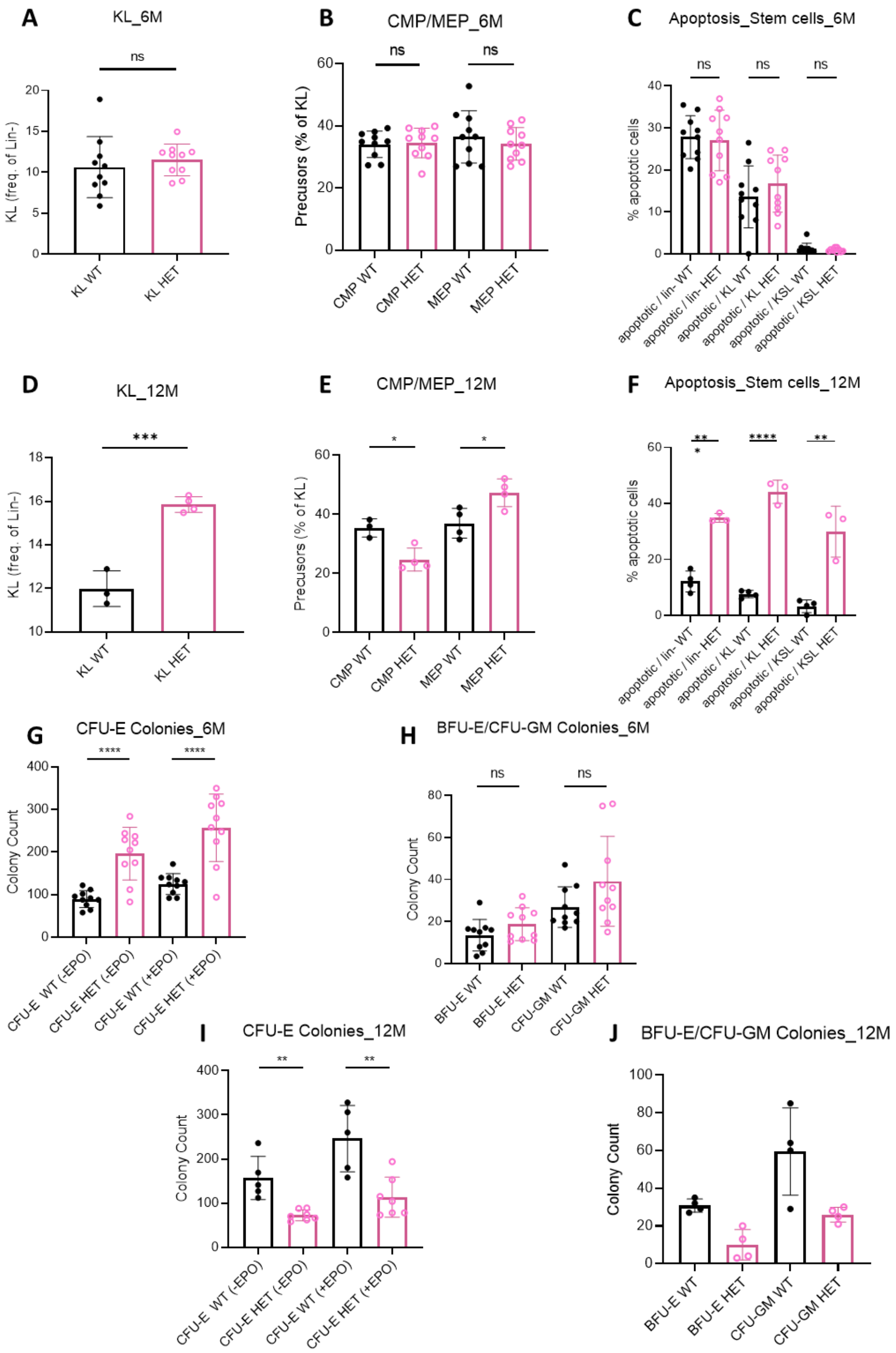


Supplemental Figure 1 (Figure S1): Frequency of Kit+/Lin- cells (A,D) in bone marrow, proportions of CMPs and MEPs (B,E), and proportions of apoptotic cells in Lin-, Kit+/Lin- and Kit+/Sca+/Lin- cells (C,F) in 6 months (A,B,C) and 12 months old (D,E,F) female animals. Colony forming units (CFU) of the erythroid lineage with and without erythropoietin stimulation (G,I), burst forming units of the erythroid lineage and CFU of the granulocyte-macrophage lineage (H,J) in six months (G,H) and 12 months old (I,J) females.**** $p<0.0001$; *** $p>0.001$, ** $p<0.01$, * $p<0.05$
